# Reactivation of transposable elements following hybridization in fission yeast

**DOI:** 10.1101/2021.08.02.454278

**Authors:** Sergio Tusso, Fang Suo, Yue Liang, Li-Lin Du, Jochen B.W. Wolf

**Author notes:** authors to whom correspondence should be addressed, Corresponding authors: Sergio Tusso, Jochen B. W. Wolf.

## Abstract

Hybridization is thought to reactivate transposable elements (TEs) that were efficiently suppressed in the genomes of the parental hosts. Here, we provide evidence for this ‘genomic shock hypothesis’ in the fission yeast *Schizosaccharomyces pombe*. In this species, two divergent lineages (*Sp* and *Sk*) have experienced recent, likely human induced, hybridization. We used long-read sequencing data to assemble genomes of 37 samples derived from 31 *S. pombe* strains spanning a wide range of ancestral admixture proportions. A comprehensive TE inventory revealed exclusive presence of long terminal repeat (LTR) retrotransposons. Sequence analysis of active full-length elements, as well as solo-LTRs, revealed a complex history of homologous recombination. Population genetic analyses of syntenic sequences placed insertion of many solo-LTRs prior to the split of the *Sp* and *Sk* lineages. Most full-length elements were inserted more recently after hybridization. With the exception of a single full-length element with signs of positive selection, both solo-LTRs, and in particular, full-length elements carried signatures of purifying selection indicating effective removal by the host. Consistent with reactivation upon hybridization, the number of full-length LTR retrotransposons, varying extensively from zero to 87 among strains, significantly increased with the degree of genomic admixture. This study gives a detailed account of global TE diversity in *S. pombe*, documents complex recombination histories within TE elements and provides evidence for the ‘genomic shock hypothesis’.

## Introduction

Hybridization is a pervasive evolutionary force with implications for adaptation and species diversification (Abbott et al. 2013). It entails disruption and novel arrangement of parental haplotypes as essential upshots with the potential to alter regulatory pathways (Turner et al. 2014). This includes regulation and epigenetic control of transposable elements (TEs) (Han et al. 2004) with proven consequences for speciation (Serrato-Capuchina and Matute 2018) and genome evolution (Kazazian 2004). Barbara McClintock hypothesized that hybridization could lead to a “genomic shock” reactivating the mobilization of TEs that were efficiently suppressed in the parental genomes (McClintock 1984). This hypothesis follows from the idea of a co-evolutionary arms race (Van Valen 1973) between TEs striving to maximize proliferation and the host genome evolving suppression mechanisms to keep TE activity in check. By introducing elements of untested genetic variation into a naïve genomic background, hybridization has the potential to disrupt genome stability with the possible effect of reactivating TEs (McClintock 1984).

Evidence for the ‘genomic shock hypothesis’ is scare, despite investigation in a diverse array of species. Results are often mixed, and outcomes differ even between closely related species. For example, intraspecific crosses between *Drosophila melanogaster* males containing the P-element transposon with naïve females lacking expression of the suppressor gene result in hybrid dysgenesis (Kidwell et al. 1977; Bingham et al. 1982; Kidwell 1983; Bucheton et al. 1984). In other species of *Drosophila* this effect cannot be consistently replicated (Coyne 1985; Hey 1988; Lozovskaya et al. 1990; Vela et al. 2014). Hybridization between *Arabidopsis thaliana* and *A. arenosa* induces up-regulation of ATHILA retrotransposon expression and reduces hybrid viability (Josefsson et al. 2006). However, such an effect is not observed in crosses between *A. thaliana* and *A. lyrata* (Göbel et al. 2018). In sunflowers, contemporary crosses between *Helianthus annuus* and *H. petiolaris* show no evidence for increased large-scale TE mobilization (Kawakami et al. 2011; Ungerer and Kawakami 2013; Renaut et al. 2014). Yet, over evolutionary timescales, sunflower hybrid species combining ancestry from the same parental species show elevated number of LTR retrotransposons, indicating a role of hybridization for TE release in the past (Ungerer et al. 2006; Staton et al. 2009; Ungerer et al. 2009). Direct evidence for TE reactivation was observed from a 232-fold increase in TE expression in hybrids of incipient whitefish species (Dion-Côté et al. 2014). In other major groups like fungi, relatively little attention has been paid to study TE reactivation in the course of hybridization. In two recent studies in *Saccharomyces* species, no evidence supporting the genomic shock hypothesis was found (Hénault et al. 2020; Smukowski Heil et al. 2020).

Detailed investigation of the ‘genomic shock hypothesis’ has long been hampered by technical difficulties of accurate TE characterization limiting studies for the most part to comparative genomics between high quality assemblies of few, evolutionary divergent species (Hoban et al. 2016; Villanueva-Cañas et al. 2017; Bourgeois and Boissinot 2019). Long-read sequencing technology opens the opportunity to characterize TE variation at the resolution of multiple individual genomes from the same species. We here study the ‘genomic shock hypothesis’ at microevolutionary resolution in the fission yeast *Schizosaccharomyces pombe*. *S. pombe* is a haploid, unicellular ascomycete fungus of the Taphrinomycotina subphylum with facultative sexual reproduction (Jeffares 2018). Recent population genetic studies have shown that all globally known strains arose by recent admixture between two divergent, ancestral lineages (described as *Sk* and *Sp*) (Tao et al. 2019; Tusso et al. 2019). These two lineages most likely diverged in Europe (*Sp*) and Asia (*Sk*) since the last glacial period. Human induced migration at the onset of intensified transcontinental trade possibly induced hybridization of these ancestral lineages ∼20–60 sexual outcrossing generations ago. Hybridization resulted in a broad range of ancestral admixture proportions predicting levels of phenotypic variation and reproductive compatibility between strains (Tusso et al. 2019).

To date, a detailed TE inventory has only been conducted for a single *S. pombe* strain (972 *h^−^*) which has been used for the assembly of the reference genome (Wood et al. 2002; Bowen 2003) and is of pure *Sp* ancestry (Tusso et al. 2019). TEs found in the reference genome are all retrotransposons (class I TEs) with long terminal repeats (LTRs), which can be grouped into several LTR families (α-i) on the basis of phylogenetic analyses (Bowen 2003). The vast majority of TE elements in the reference genome only occurs in the form of solo-LTRs (174 out of 187 TE elements). Merely two types of full-length retrotransposons, called Tf1 and Tf2, containing the internal coding region (hereafter referred to as full-length elements) are known to exist in *S. pombe* (Levin et al. 1990; Levin 1995). Both Tf1 and Tf2 belong to the Ty3/Gypsy type of LTR retrotransposons, and their LTRs belong to the α and β family, respectively. In the reference genome, full-length elements (13 of 187 TE elements) are all Tf2 elements (Esnault and Levin 2015), but full-length Tf1 elements are known to exist in several wild strains (Levin et al. 1990). Short-read genome sequencing data have been used to investigate variation of TE insertions in *S. pombe* (Jeffares et al. 2015), but TE sequences cannot be reliably inferred from short reads.

In this study, using long-read sequencing data from strains spanning the world-wide diversity of fission yeast, we present a comprehensive description of the TE repertoire and place it in the context of recent hybridization between the *Sp* and *Sk* ancestors. Detailed phylogenetic and population genetic analyses were used to shed light on TE selection dynamics, assess the evidence of complex recombination and test the genomic shock hypothesis.

## Results

A previous large-scale analysis of global genetic diversity of *S. pombe* using 161 strains has identified 57 clades differing by at least 1,900 SNPs (Jeffares et al. 2015). We generated long-read sequencing data from a subset of 37 samples representing 29 of the 57 clades, and two additional, previously undescribed strains (**Figure 1, Supplementary Figure 1, Supplementary table 1**). In six instances, two clones of the same strain accessed from different labs (with potentially different recent history) were independently sequenced. The data set was further complemented with the reference genome. Analyses of SNP variation place these 38 samples well within the global continuum of *Sp* to *Sk* ancestry (Tusso et al. 2019) (**Figure 1A,B** and **Supplementary Figure 1**). Consistent ancestry profiles between clones of the same strain reflect high technical replicability (**Supplementary Figure 1** and **Supplementary Figure 2**).

**Figure 1.**
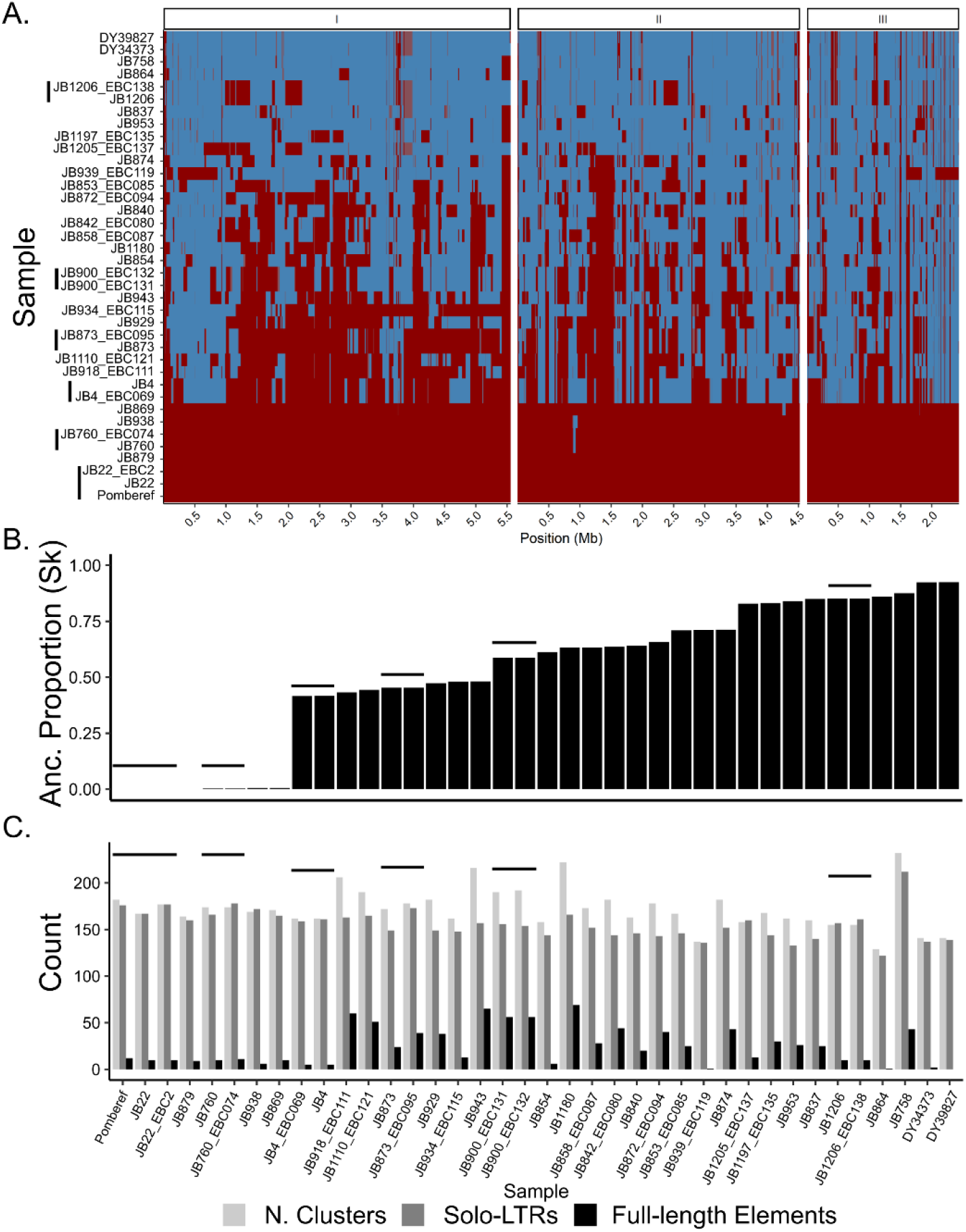
Genome composition by ancestry and LTR repertoire of a global collection of 38 samples corresponding to 31 non-clonal haploid *S. pombe* strains. A. Heatmap representing SNP-based haplotypes across all three chromosomes (I, II, III) for the 38 samples used in this study. Six clonal samples derived from the same strain are indicated by vertical bars. Haplotypes are painted by *Sp* and *Sk* ancestry shown in red and blue colour, respectively. For details on the inference of ancestry components, we refer to Tusso et al. (2019) B. Distribution of *Sk* ancestry proportion per sample. C. Number of solo-LTR sequences, full-length elements and total number of clusters per strain. For a summary of the total number of TEs per strain see **Supplementary Figure 5**. In all panels, clonal samples are grouped by horizontal black lines.

### Global TE diversity of S. pombe

From long-read data averaging 85× sequence coverage per sample (range: 40×-140×), we generated 37 individual (near-)chromosome genome assemblies (**Supplementary table 3**). Subsequent annotation allowed for characterization of TE repertoires for each individual assembly. To establish synteny of TEs between the often highly rearranged genomes (Brown et al. 2011; Tusso et al. 2019), we translated the coordinates of TEs in each individual assembly to those in the reference genome. Locations of TE elements were highly collinear between samples (**Supplementary Figure 3**) and often hosted multiple copies of TEs (as many as 16 copies). We refer to these local TE aggregations as TE clusters throughout. Within a cluster, synteny could not be unequivocally defined, restricting several subsequent analyses to TE clusters, rather than individual TE sequences.

Across all samples we identified 8,505 TE sequences that were contained in 719 TE clusters. Consistent with previous work, all TEs belonged to the Ty3/Gypsy LTR retrotransposon superfamily (Levin 1995; Levin et al. 1990). Additional identified helitron and TIR transposons overlapped with annotated genes in the reference genome and require further validation to guard against false positive inference. The vast majority of TE sequences occurred as solo-LTRs, and only 1,169 TE sequences were longer than 1.5 kb and contained internal sequences. The vast majority of the long sequences (924 sequences) had a length of around 4.9 kb, the expected length of full-length elements (**Figure 1C**, **Supplementary Figure 4**). The number of TE elements varied between strains both for solo-LTR sequences (range 115-198), as well as for full-length elements (range: 0 - 67, **Figure 1C, Supplementary Figure 5**).

### Methodological comparison

Variation between clones derived from the same strain was small, but non-zero for the total number of clusters and TE sequences. A fraction of the differences was likely not owing to genome quality differences, but may reflect biological variation acquired during a short period of time. (**Supplementary Figure 6**).

The overall high consistency of TE sequences in our clonal *de-novo* genomes contrasted with low congruence with TE inference from other sources (**Supplementary Figure 7**). A comparison between our annotation of TE sequences in the reference genome and a BLAST-based annotation of TE sequences in an earlier version of the reference genome reported by Bowen *et al*. (2003) (**Supplementary table 6**) showed consistency in 117 identified clusters. 47 and 13 clusters showed unique evidence in either our Pomberef annotation or Bowen’s annotation. Additionally, comparing presence/absence of TE clusters as inferred from short-read data (Jeffares et al. 2015) (**Supplementary Figure 7**) to our annotations revealed a large proportion of inconsistent clusters ranging from 23% (JB22) to 49% (JB874). In summary, these results highlight the limitation of short-read data to infer TE insertions, confirm the robustness of long-read based inference and tentatively suggest rapid variation between clonal strains.

### LTR diversity is superimposed on ancestral population divergence

Next, we extracted solo-LTRs and flanking LTR sequences from full-length elements (both 5′ and 3′) amounting to 9,503 LTR sequences altogether. Phylogenetic analysis of the resulting sequence alignment provided an overview of the global diversity of LTR sequences in *S. pombe* (**Figure 2A**). Although bootstrap support was generally low, LTR sequences could be broadly grouped into previously reported families (Bowen 2003). In most families, LTRs occurred exclusively as solo-LTRs, had long terminal branches and showed high intra-family diversity. This phylogenetic signature reflects past transpositions of now extinct elements, and is consistent with a long history of recombination-mediated conversion of full-length elements into solo-LTRs followed by pseudogenization of the remaining LTR sequences. The α and β families constitute an exception. These two families hosted the large majority of full-length elements found in the data set, Tf1 and Tf2, respectively (**Figure 2B**). The other group of LTR sequences associated with full-length elements, here coined *Sub-α/ζ*, was closely related to the α family, but showed evidence of recombination between LTR haplotypes from the α and β families, or between the β family and an ancestral sequence related to the ζ family (**Supplementary Figure 8**). In total, we identified at least 24 recombinant solo-LTR haplotypes, several of which were found in multiple clusters (up to 64 and 40 clusters for the two most common recombinant haplotypes) (**Supplementary Figure 9**).

**Figure 2.**
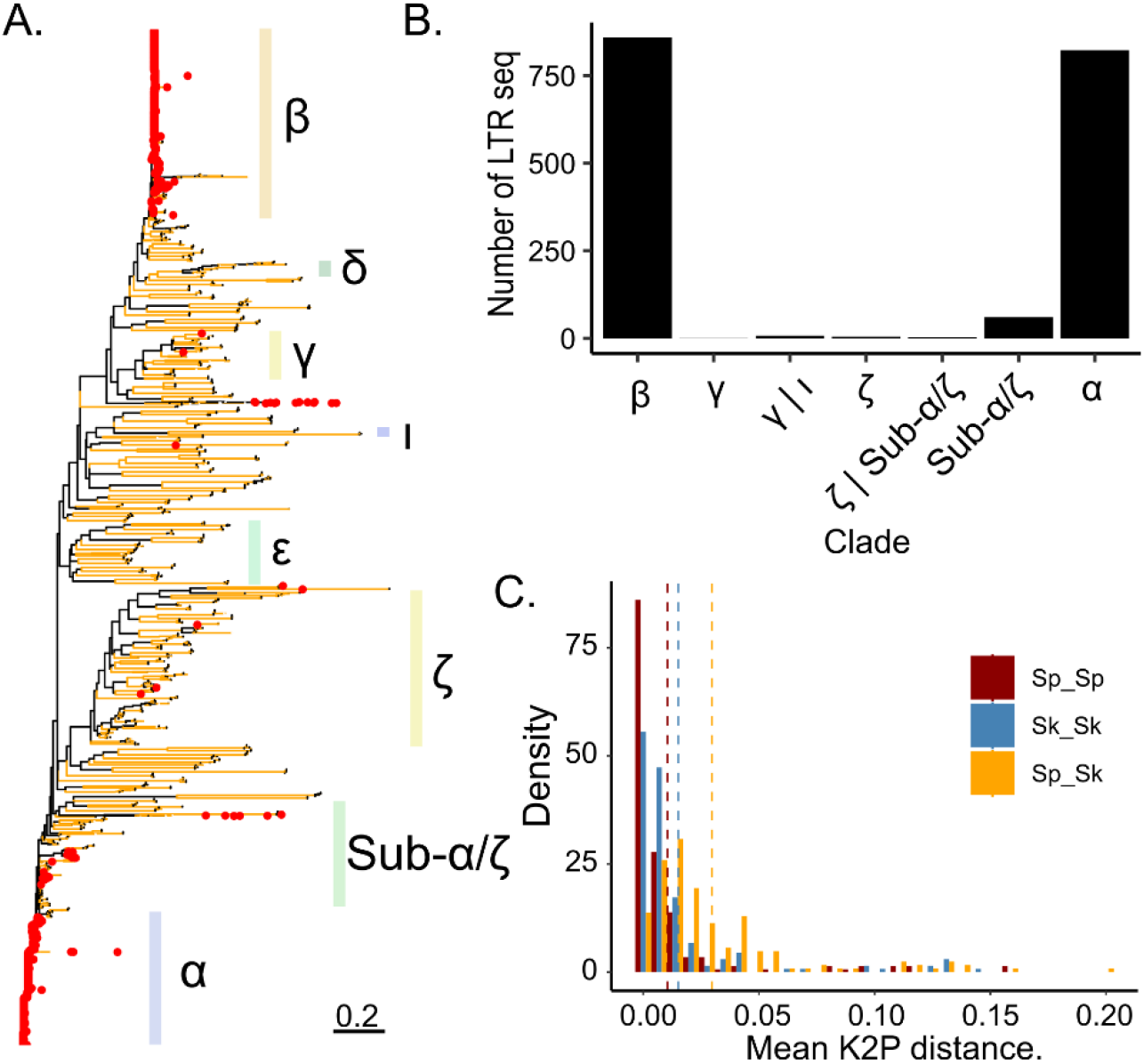
Phylogenetic relationship between LTR sequences. A. Maximum likelihood un-rooted tree for solo-LTRs and LTRs flanking full-length elements. Branches with bootstrap support higher than 95 are shown in yellow. LTR sequences associated with full-length elements are indicated with red points. Nomenclature of families follows Bowen et al. (2003) (see methods). **B.** Number of flanking LTRs from full-length elements grouped by LTR family. Sequences without clear family membership, are classified by the two most closely related families. **C.** Pairwise divergence of LTR sequences belonging to the same family within syntenic clusters. Divergence between sequences is grouped by ancestral background: within ancestral background (*Sp vs. Sp* or *Sk vs. Sk*) or between ancestral backgrounds (*Sp vs. Sk*). Mean divergence per group is indicated by hashed, vertical lines.

To relate the diversity of LTR sequences to global species diversity (*Sp* vs. *Sk* lineage) we inferred the ancestral genomic background of each syntenic cluster for each strain. In 581 of all 662 syntenic clusters, LTR sequences from the same family were exclusively present in clusters embedded by either of the two ancestral backgrounds. Consistent with higher overall genetic diversity in the *Sk* lineage (Tusso et al. 2019), the percentage of clusters with lineage-specific LTR insertions was higher for the *Sk* than for the *Sp* background (57% and 37% respectively). In 152 clusters, sequences originating from the same LTR family occurred in at least one sample of each ancestral background, in 113 in at least two. Co-occurrence across ancestral backgrounds makes these sequences prime candidates of ancestral insertions prior to divergence of the two lineages. Across all shared 113 syntenic clusters, mean pairwise divergence was lower between sequences from the same LTR family inserted into the same ancestral background (*Sp-Sp* and *Sk-Sk*) than between different ancestral backgrounds (*Sp-Sk*) (**Figure 2C** and **Supplementary Figure 10**). Moreover, mean pairwise distance within the *Sk* group was higher than for the *Sp* group, which is consistent with higher effective population size inferred for the *Sk* group (Tusso et al. 2019). In summary, these results from solo-LTR sequences are consistent with the two-clade history inferred from genome wide SNPs (Tao et al. 2019; Tusso et al. 2019) and show that a proportion of solo-LTRs preceded *Sp* and *Sk* divergence and subsequent hybridization. This includes LTR sequences from the two most common α and β families that are characteristic of full-length elements (**Supplementary Figure 11**).

### Haplotype diversity of full-length elements documents a history of recombination

Next, we focused on full-length elements for which two haplotypes, Tf1 and Tf2 elements, have been previously described. Using window-based haplotype painting, all sequences were collapsed into 11 discrete haplotypes, each present in at least 5 sequences (see Methods, **Figure 3A**). These haplotypes can similarly be identified by means of phylogenetic analyses (**Figure 3B**). Differentiation between haplotypes was primarily due to divergence in the flanking LTR sequences and the first ∼2 kb of the internal sequence. Prevalence in the data set was highest for the Tf1 and Tf2 haplotypes occurring in 207 and 203 clusters, respectively. The remaining 9 haplotypes populated 69 clusters. Despite relatively lower numbers, paralogous occurrence across different genomic regions (clusters) suggests that some of these recombinant haplotypes have recently been actively transposed. Prominent examples are haplotypes Tf2e, Tf2f and Tf2g found in 6, 9 and 28 independent clusters, respectively.

**Figure 3.**
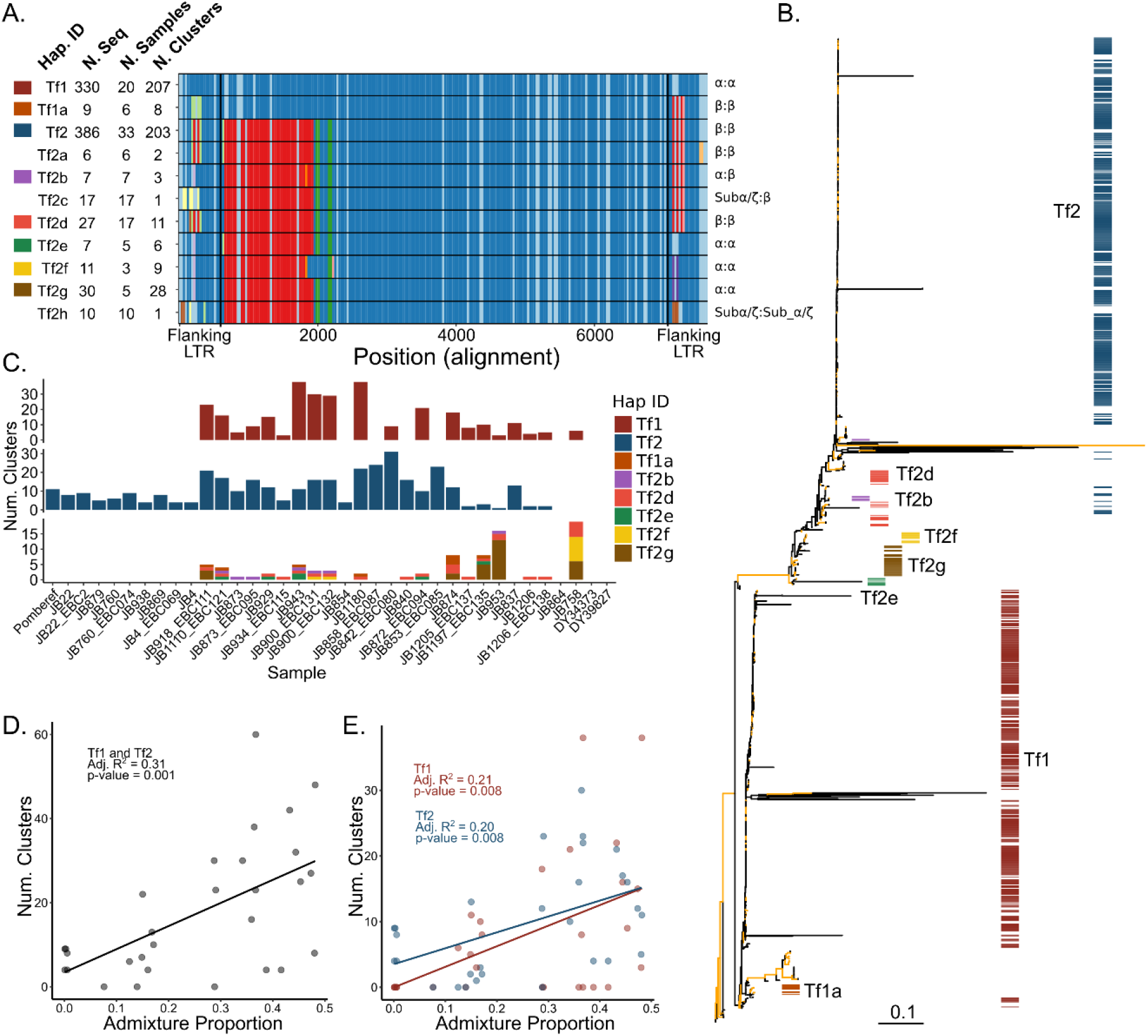
Diversity of full-length LTR elements. a. Alignment of the 11 haplotypes identified by window-based haplotype painting in a global sample of *S. pombe*. For each haplotype we show: haplotype ID, number of sequences found in all samples, number of samples, and number of independent clusters. Vertical black lines show boundaries of flanking LTRs. Colours per window (vertical comparison within the alignment) represent haplotype difference from TF1 used as reference. Note that due to insertion-deletion polymorphism the alignment exceeds the length of the individual full-length LTR elements. Flanking LTR families per haplotypes are shown on the right (5’LTR:3’LTR). **b.** Maximum likelihood un-rooted tree for full-length LTRs. Branches with bootstrap support higher than 95 are shown in yellow. Colours correspond to the colour assignment of the eight most common haplotypes in panel a. **c.** Number of common haplotypes for full-length elements found in at least 3 independent clusters shown per sample (lower panel). Haplotypes Tf1 and Tf2 are shown in independent plots (upper, middle panel). Colours per haplotype ID as indicated. Samples are ordered by ancestral admixture from pure *Sp* to pure *Sk* as in Figure 1. **d.** Relationship between ancestral *Sp* and *Sk* admixture proportions and the number of clusters with Tf1 and Tf2 full-length elements. Each point represents a non-clonal strain. The adjusted proportion of total variance explained (R^2^) and the type 1 error probability (p-value) are shown as inset. **e.** As panel d, but differentiating between haplotypes TF1 and TF2.

Haplotype diversity was larger for derivates from Tf2 (9 haplotypes) than for Tf1 with only one additional haplotype. Haplotype diversity was mostly governed by homologous recombination between the Tf1 and Tf2 haplotypes. For example, haplotype Tf1a contains an internal sequence of the Tf1 haplotype, but flanking LTRs are more similar to those of Tf2 (β LTR family). Conversely, haplotypes Tf2e and Tf2g are most similar to Tf2 in the internal sequence, but its flanking LTRs are more related to those found in Tf1 (α LTR family). Other haplotypes also suggest recombination of the internal sequence, as is illustrated in Tf2f.

### Support for the genomic shock hypothesis

To estimate the age of insertion, we calculated pairwise divergence between the 5′ and 3′ flanking LTR sequence for each full-length element. Naturally, divergence in recombinant full-length elements was elevated with values exceeding 10 % (**Supplementary Figure 12**). For the vast majority of non-recombinant full-length elements of both Tf1 and Tf2, however, divergence was consistently lower than 1%, suggesting that TEs mobilized over a short period of time in a ‘burst of activity’ rather than accumulated gradually. Recent activity succeeding the *Sp* and *Sk* split was further supported by an uneven segregation of full-length haplotypes between ancestral backgrounds: the Tf2 haplotype was found in the majority of samples, but was absent in strains with predominant *Sk* ancestry. The Tf1 haplotype showed the reverse pattern (**Figure 3C**). Other haplotypes were either sample specific or were restricted to a few samples, with JB758 and JB953 being exceptionally prolific hosts of the non-typical full-length haplotypes. Haplotypes whose prevalence was restricted to only a few samples often showed high abundance within those samples. For instance, Tf2f and Tf2g were restricted to 3 and 5 samples, but populated 9 and 28 clusters, respectively.

Overall, this analysis suggests that full-length haplotypes can be grouped into two classes. i) Two common haplotypes (Tf1 and Tf2) which are characterized by flanking LTRs of the α and β family and are found in most samples, but segregate at different rates in the ancestral groups (with dominance of Tf2 in *Sp* and dominance of Tf1 in *Sk*). Analysis of solo-LTRs suggest likely presence of at least the β family prior to the split of *Sp* and *Sk* (**Supplementary Figure 11**). ii) A number of haplotypes restricted to few strains, often with evidence for recombination. Based on sequence similarity, at least some of these rarer haplotypes originated by homologous recombination between Tf1 and Tf2 and remained active thereafter.

We next examined whether the patterns of TE diversity in *S. pombe* conform to the ‘genomic shock hypothesis’. If recent hybridization (∼20-60 sexual generations ago, Tusso et al. 2019) reactivated TE activity, strains with admixed genomic backgrounds should, on average, host a larger number of full-length elements relative to non-admixed strains. This prediction was supported by the data. We observed a significant positive relationship between ancestral admixture proportion and the number of clusters containing full-length Tf1 or Tf2 sequences (including only non-clonal samples; **Figure 3D**). This correlation remained if each haplotype was considered independently (**Figure 3E**), and when exclusively focusing on the most recent full-length element insertions (singleton clusters; p-value 0.004, adj. R^2^: 0.24; Supplementary Figure 1**3**). This correlation is consistent with increased TE activity in admixed samples. Alternatively, it may reflect a demographic signal of an increased population mutation rate in a pool of hybrids having recently experienced a population expansion. Under this scenario, we would expect to observe a general excess of low frequency variants not only in TEs, but any type of genetic variation. Repeating the analysis using neutrally evolving SNPs did, however, not support a differential demographic explanation. There was no indication that the number of singleton SNPs was elevated in admixed samples (p-value 0.088, adj. R^2^: 0.07; Supplementary Figure 1**3**). Moreover, no correlation was found between the number of singleton SNPs and number of singleton TE clusters (p-value 0.374, adj. R^2^: -0.01; Supplementary Figure 1**3**). On the basis of these results, we propose that recent hybridization increased the rate of TE proliferation in *S. pombe*, as is predicted by the ‘genomic shock hypothesis’.

### Population genetic inference of selection

To shed further light on the evolutionary history of TE elements in *S. pombe*, we constructed unfolded site-frequency spectra (SFS) scoring presence/absence of clusters as allelic states. First, we considered all clusters found in non-clonal strains (686 in total). 508 clusters (74.0%) were found in no more than five samples, and 363 (52.9% of total) were restricted to single samples (singletons) (**Figure 4A**). 115 clusters (16.7%) occurred in 90% or more of the samples. These ubiquitously present clusters contained predominantly solo-LTRs.

**Figure 4.**
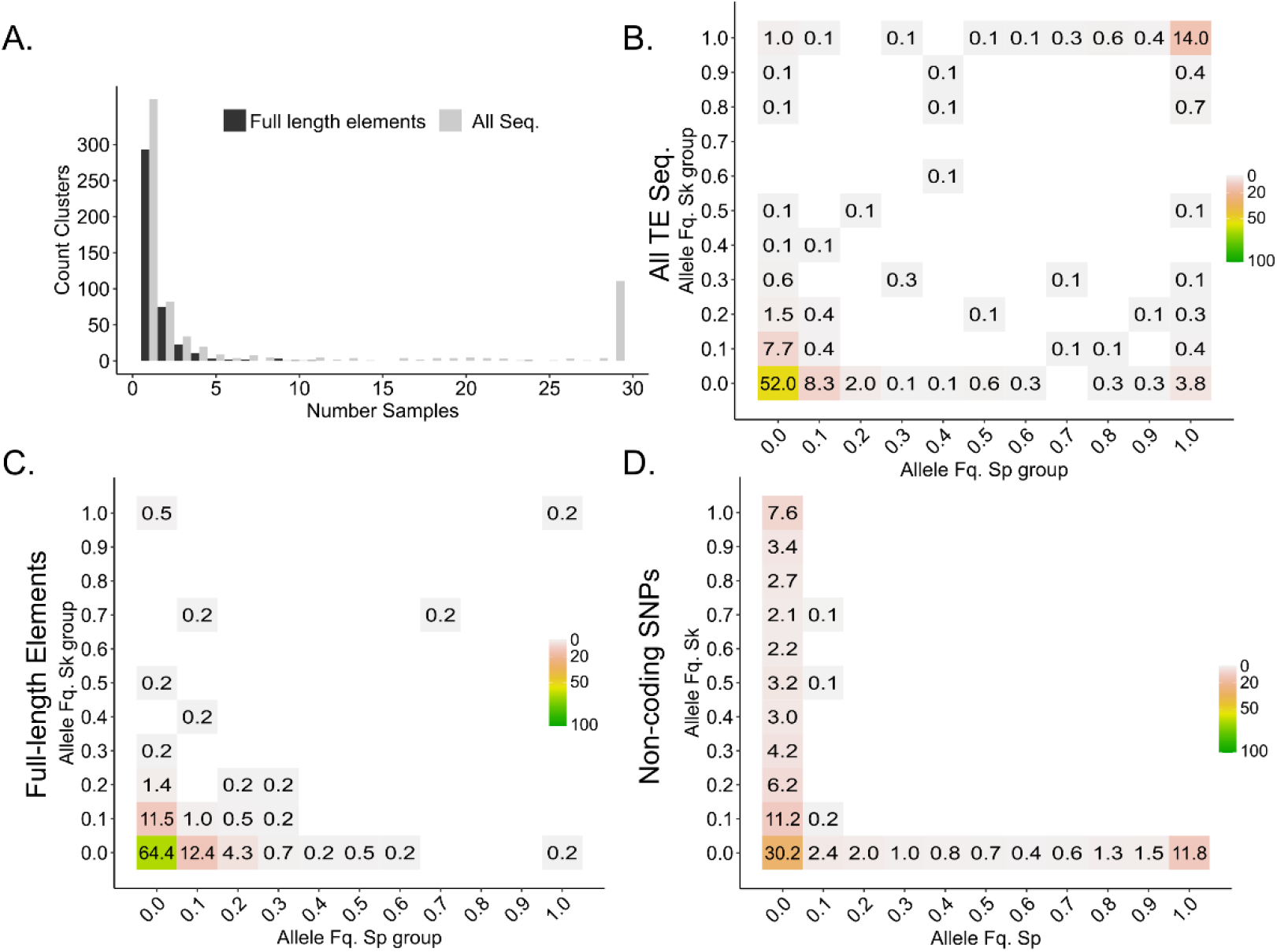
**Allelic variation of LTR clusters between non-clonal strains**. **A.** One-dimensional site frequency spectrum summarizing the allelic frequency of TE cluster insertions across all 31 non-clonal strains. Bars differentiate between either all sequences or full-length elements. B, C, D. Two-dimensional site frequency spectra showing the proportion of allele frequency bins shared between the ancestral *Sp* and *Sk* background for all TE sequences **(B)**, full-length elements **(C)** and non-coding genome-wide SNPs **(D)**. Note that site frequency spectra in B and C are unfolded, but folded in d. For folded spectra of B and C see **Supplementary Figure 14**. Number and colour range indicate the percentage out of all variants within each panel.

Restricting the SFS to to full-length elements, low frequency variants were substantially more common and high frequency clusters were drastically reduced. Of 415 clusters with at least one full-length element, 423 (97.4%) were found in no more than five samples and 258 (59.4% of total) were singletons. Only two full-length element-containing cluster exceeded a frequency of 30%.

The excess of low-frequency and depletion of high-frequency clusters containing full-length elements can result from three non-mutually exclusive processes: a recent burst in transposition rate, a recent demographic population expansion resulting in an increase of rare variants and purifying selection removing new insertions and maintaining variants in low frequency. To evaluate the effect of demographic variation, the history of admixture of the species has to be accounted for. We therefore scored variation of SNPs adjacent to each cluster and inferred the ancestral origin for each cluster (*Sp* or *Sk*). Subsequently, we calculated the two-dimensional site frequency spectrum (2dSFS) for both *Sp* and *Sk* ancestry using only non-clonal samples (**Figure 4B**). Considering all TE elements (solo-LTR and full-length), ∼15% of the clusters were fixed in both ancestral lineages likely representing ancestral insertions present prior to the split of the two lineages. The majority of clusters, however, were lineage-specific and in low frequency, with 43% of the clusters having frequencies below 0.1 in both lineages (∼68% in folded SFS – **Supplementary Figure 14**). For full-length elements, low frequency variants were substantially more common (**Figure 4C** and folded 2dSFS in **Supplementary Figure 14**). Here, the percentage of low frequency variants increased to ∼64% and clusters at intermediate frequency were reduced. To relate these patterns to genetic variation presumably evolving neutrally, we constructed the folded 2dSFS of genome-wide non-coding SNPs (**Figure 4D**). Here, the percentage of low frequency variants for non-coding SNPs was ∼30% which was below values of the folded 2dSFS from all-LTR (67.9%) and full-length variants (64.6%) (**Supplementary Figure 14).** The relative increase of rare alleles in LTRs over neutral SNPs cannot be explained with demographic expansion alone. Instead, these results are best explained by a recent increase in TE proliferation (in admixed genomes, see above) and a likely additional component of purifying selection against LTR elements in general. Consistent with the latter prediction, the percentage of fixed clusters was higher in the *Sp* ancestral population where a lower *Ne* has been predicted (Tusso et al. 2019) reducing the efficacy of selection (Charlesworth and Charlesworth 1983; Charlesworth and Langley 1989).

## Discussion

### Support for the genomic shock hypothesis on a microevolutionary scale

This study provides empirical evidence for a role of hybridization in the evolution of transposable elements. It adds to the evidence of TE reactivation as a consequence of hybridization in fungi (cf. Hénault et al. 2020; Smukowski Heil et al. 2020). Our results are consistent with the idea that hybridization of two closely related *S. pombe* lineages, *Sp* and *Sk* (D_xy_ ∼ 0.005), activated TE proliferation in the admixed genomes (‘genomic shock’ (McClintock 1984)). The recent timing of hybridization allowed us to witness the sudden burst of transposition in natural populations, before signal loss by re-establishment of novel repression mechanisms. Sudden bursts of transposition generated a large cohort of insertions with roughly the same age, as has similarly been documented in *Drosophila* (Vieira et al. 1999), rice (Piegu et al. 2006), piciformes (Manthey et al. 2018), salmonids (de Boer et al. 2007), and primates (Pace and Feschotte 2007).

Hybridization may have contributed to increased numbers of full-length TEs in admixed genomes in three ways. First, by import of active, full-length elements from one pure parental background into the other. This was illustrated by transfer of Tf2 full-length elements from the *Sp* genomic background into the originally Tf2-free *Sk* background in admixed samples. Second, by disruption of the allelic inventory of co-adapted control mechanisms impeding TE mobilization in the parental backgrounds (e.g. CENP-B proteins (Cam et al. 2008), Set1 histone methyltransferase (Lorenz et al. 2012), histone deacetylases and chaperones (Murton et al. 2016), and to a low degree the RNAi machinery (Woolcock et al. 2011; Chapman 2018)). While the precise molecular mechanisms underlying activity, repression and copy number of TEs have been extensively studied in the reference strain of *S. pombe* (Hansen et al. 2005; Cam et al. 2008; Lorenz et al. 2012; Murton et al. 2016), the mechanism underlying the observed reactivation of TEs in natural populations remains elusive and will require further study.

A third way of how hybridization may have contributed to TE proliferation is by reduction of *Ne* in the founder population of hybrids, reducing efficiency of purifying selection against TE (Charlesworth 2009; Charlesworth and Charlesworth 1983). While a bottleneck as recent as 20 outcrossing generations ago is difficult to reconstruct, patterns of TE frequency overall confirmed the relationship between effective population size and purifying selection. Contrary to findings of other hybridizing yeast lineages, we found that putatively active full-length elements are only ubiquitously present in one of the pure parental backgrounds (*Sp*). Strains predominated by *Sk* ancestry had an overall reduced number of solo-LTRs, less fixed LTR insertions, and hosted fewer, if any, copies of full-length LTR elements (with only few, slightly admixed strains hosting Tf1). More efficient TE control and removal in the *Sk* lineage is consistent with its higher effective population size predicting higher efficiency of purifying selection of deleterious TE elements compared to the *Sp* ancestral lineages (Lynch and Walsh 2007). Similar variation in TE content as a result of changes in *Ne* has been observed in *Brachypodium* (Stritt et al. 2018), *Arabidopsis* (Lockton et al. 2008), *Caenorhabditis* (Dolgin et al. 2008), *Drosophila* (García Guerreiro et al. 2008), sticklebacks (Blass et al. 2012), *Anoles* (Tollis and Boissinot 2013), humans and mice (Xue et al. 2018).

Transposition-selection balance was further reflected by site frequency spectra skewed towards low frequency of TE insertions (Barrón et al. 2014; Bourgeois and Boissinot 2019). Analysis in *Drosophila* found that 48 to 76% of TEs had frequencies lower than expected under neutrality as a consequence of purifying selection (Cridland et al. 2013; Barrón et al. 2014; Blumenstiel et al. 2014). Deviation from neutral expectations has also been observed in other systems like *Anoles* (Ruggiero et al. 2017), mice (Xue et al. 2018), or *Arabidopsis* (Hazzouri et al. 2008; Lockton et al. 2008). In this study, we similarly found 68.4% (85.6% in folded 2dSFS) of TE insertion in frequencies below 0.2, contrasting with 44.1% found for presumably neutral, non-coding SNPs.

Overall, the data are best explained by a burst in transposition rate upon hybridization on a background of continuous purging of both solo-LTRs, and to a larger degree full-length elements. To quantify the contribution of purifying selection on TE expansion during and after hybridization explicit demographic reconstruction and fitness data would be necessary.

### TE dynamics through time – the role of homologous recombination

The combination of phylogenetic and population genetic analyses of this study further contributes to our understanding of TE dynamics in natural populations. Under expectations of the neutral evolutionary theory, younger TE insertions are on an average expected to segregate at lower allele frequencies than older insertions (Kimura and Ohta 1973). Moreover, under the assumption of a molecular clock, progressively older insertions will have accumulated increasingly more mutations (Petrov et al. 1996). Thus, both allele frequency and sequence divergence provide information on the age of TE insertions (Blumenstiel et al. 2014). Across our global sample of *S. pombe* strains, the vast majority of full-length LTR elements segregated at very low frequencies, and sequence divergence of flanking LTRs within and between copies was shallow. In contrast, solo-LTRs segregated at higher frequencies and were diversified into multiple, divergent families. These results are consistent with full-length elements being mostly of young age and solo-LTRs being of much older origin.

In the *S. pombe* reference strain, ∼70% of Tf2 mobilization events involve homologous recombination between newly synthesized cDNA and a pre-existing copy of Tf2; the remaining ∼30% of cases integrate in novel chromosomal locations (Hoff et al. 1998). Homologous recombination and recycling of target sites has two consequences for TE dynamics. First, older TE insertions may be replaced by more recent events, hampering the reconstruction of TE dynamics over long evolutionary time scales. Second, homologous recombination can be an important mechanism for TE diversification, resulting in new recombinant haplotypes for which this study provides copious examples. Importantly, recombination has occurred between elements of divergent families (Tf1 and Tf2) generating chimeric elements. Several of these recombinant haplotypes were found in multiple genomic clusters suggesting they have been recently active. Although most chimeric elements contained identical flanking LTRs, this was not always the case. In several instance, LTRs differed between the 3’ and 5’ end (cf. **Supplementary Figure 12**). However, with the only exception of haplotype Tf2b found in 3 clusters, all other haplotypes were found in single clusters. Since flanking LTR are known to be homogenized during reverse transcription, chimeric elements with divergent flanking LTR are more likely the result of homologous recombination between cDNA and a previous TE insertion.

### Adaptive evolution of TEs

Transposable elements do not exclusively cause damage to the host. As sources of molecular variation, they can reroute regulation of gene expression (Sundaram and Wysocka 2020; Trizzino et al. 2017) and contribute to adaptive evolution (Schrader and Schmitz 2019). Environmental change can influence transposition rate as observed in *S. cerevisiae* (Paquin and Williamson 1984), or provide opportunities for positive selection of novel TE insertions (Aminetzach et al. 2005; Gresham et al. 2008; Hof et al. 2016; Esnault et al. 2019). Over long evolutionary time scales, beneficial TE insertions can become domesticated by the host genome (Miller et al. 1999). In *S. pombe*, for instance, CENP-B proteins involved in TE silencing (Cam et al. 2008) are believed to have evolved from a domesticated pogo-like DNA transposase (Irelan et al. 2001; Casola et al. 2008).

TE insertions in *S. pombe* have also been discussed in the context of adaptation to environmental disturbance, or stress response. TE expression has been shown to be induced under stress conditions (Chen et al. 2003; Sehgal et al. 2007), and an enhancer sequence contained in Tf1 induces expression in adjacent genes (Leem et al. 2008). Artificially induced Tf1 insertions in the reference strain preferentially occurred upstream of stress response genes (Guo and Levin 2010). In natural strains, TE insertions are also enriched in the proximity to promoters of gene for stress response (Jeffares et al. 2015), which has been interpreted as evidence for TE induced adaptive response (Esnault et al. 2019). Additionally, analysis of experimental populations under different environments shows variation in the genomic distribution of Tf1 integrations, as well as increased transposition rates under stress conditions (Esnault et al. 2019). Competition assays of the same evolved populations showed a selective advantage for several TE insertions under stress conditions with heavy metals.

In the context of this study, it is conceivable that the human-associated, hybrid strains that have been rapidly dispersed across the globe experienced a range of novel, suboptimal environments inducing stress response (Jeffares 2018). Indeed, most strains used in this study were isolated from diverse substrates (Jeffares et al. 2015). Despite the apparent opportunities for adaptive evolution, the vast majority of full-length elements segregated at lower than expected frequency. This suggests that most of them were inserted recently (transposition burst) and rapidly removed by purifying selection and/or reduced to solo-LTRs by homologous recombination, limiting the time frame of active proliferation. Our results do not exclude the adaptive potential of TEs, but instead suggest limitation of adaptive evolution to short periods of stress after which the selective advantage is lost. In this study, we identified only one full-length element candidate for pervasive long-term positive selection present in both ancestral backgrounds. However, the functional significance of this insertion is not clear and warrants future experimental exploration.

This study offers a comprehensive characterization of the global diversity of transposable elements in *S. pombe*. Phylogenetic and population genetic approaches provide evidence for the ‘genomic shock hypothesis’ in fungi and copious examples of homologous recombination among full-length and solo-LTR sequences. Consistent with established principles of molecular evolution, TE insertions were generally subject to purging with the exception of a single locus. These results contribute to the debate on the role of TEs in evolution, notably in speciation and adaptation (Serrato-Capuchina and Matute 2018).

## Methods

Previously, a world-wide collection of 161 naturally occurring *S. pombe* strains (JB strains) has been grouped into 57 clades differing by at least 1,900 SNPs (Jeffares et al. 2015). Within each clade, strains are near clonal. In this study, we compiled single-molecule long-read sequencing data for strains representing 29 clades that cover the spectrum of genetic variation in the species by i) selecting samples along the phylogeny (**Supplementary Figure 2**), ii) including previously reported genetic groups (Jeffares et al. 2015; Tusso et al. 2019) and iii) considering genomic variation of strain ancestry (**Figure 1** and **Supplementary Figure 1**). For 17 strains that correspond to 16 clades, data are publicly available (Tusso et al. 2019). Two of the 17 strains, previously referred to as EBC131_JB1171 and EBC132_JB1174, were found to belong to the same clade represented by the strain JB900, and were thus referred to as JB900_EBC131 and JB900_EBC132 in this study. Another set of single-molecule long-read sequencing data from 20 strains were generated in this study. These 20 strains include strains covering 13 additional clades, two strains not belonging to the 57 clades, and 5 strains sharing cluster affiliation with 5 of the 17 previously published strains. With the inclusion of the already assembled reference genome, the final data set was comprised of 38 samples (**Supplementary table 1**).

### Genome assemblies

Additional to previously published genome assemblies, PacBio long-read sequencing data were generated for the other 20 strains. We performed *de novo* assembly using two assemblers: Canu 1.8 and wtdbg 2.4 (Koren et al. 2017; Ruan and Li 2020). The parameters settings were ’genomeSize=12.5m’ for Canu and ’-x sq -L 3000 -g 12.5m’ for wtdbg. For JB4 and JB1180, which were sequenced using the PacBio RS II platform, we used the option ’-x rs’ when running wtdbg. QUAST 5.0.2 was used to evaluate the assembly quality (Gurevich et al. 2013) . For each strain, only the assembly with the superior quality was kept for further improvement. GCpp 0.0.1 was run to polish the assemblies using the Arrow algorithm and long reads. Subsequently, finisherSC 2.1 was applied to further improve the assembly (Lam et al. 2015), followed by another round of GCpp polishing. For JB1180, the assemblies generated were of poor quality, and we instead used SMRT Analysis Software 2.3.0. This pipeline resulted in general near-chromosome level genomes with a median number of contigs of 8, average N50, total assembly length, and recovery of annotated genes of 2.48 Mbp, 12.56 Mbp (reference genome 12.55 Mbp), and 97.0% respectively (**Supplementary table 2).**

### Phylogenetic analyses and the inference of ancestry blocks

We performed two analyses using SNP variants: first, phylogenetic analyses using genome-wide SNP data (Supplementary Figure 2); and second, a reported pipeline to identify the composition of *Sp* and *Sk* ancestral haplotype blocks along the genome (Supplementary Figure 1) (Tusso et al. 2019). For SNPs derived from short-read sequencing data, we used a publicly available data set in variant call format (VCF) (Jeffares et al. 2015). SNP variation of the *de-novo* genomes (this study) was inferred via alignment to the reference genome (ASM294v264) (Wood et al. 2002) and subsequent characterization of variants with the package MUMmer 3.23 (function *show-snps*) (Kurtz et al. 2004). Repetitive sequences in the reference were identified with RepeatMasker 4.0.8 (Smit et al. 2013) and excluded for SNP variant calling. For the phylogenetic analyses, genome sequences were reconstructed by editing the reference genome with the SNP information for each sample using a customized Python script (Van Rossum and Drake 2009). The analysis was performed independently for each chromosome. Alignments of all samples, including short and long read data, were used to build a maximum likelihood tree using RAxML 8.2.10-gcc-mpi (Stamatakis 2014) with default parameters, GTRGAMMAI approximation, final optimization with GTR + GAMMA + I and 1,000 bootstraps.

For inference of ancestral *Sp* and *Sk* haplotype blocks along the genome we followed (Tusso *et, al.* 2019). Strain identities and ancestral block distribution were consistent between short-read data (Jeffares et al. 2015) and long-read data (this study).

### Genome annotation of transposable elements

For each *de-novo* genome, we followed the CARP wrapper (Zeng et al. 2018) to identify and annotate repetitive sequences. Each family was annotated, and TEs were identified from other repeat sequences using a reference of known TE sequences for *S. pombe* and other fungi, obtained from the database Repbase (Bowen 2003; Bao et al. 2015). Unidentified sequences were compared to protein sequences, transposable elements in other species and retrovirus sequences using BLAST 2.7.1. Sequence references were obtained from the data base hosted by the National Center for Biotechnology Information (https://www.ncbi.nlm.nih.gov/) using the following search terms: reverse transcriptase, transposon, repetitive element, RNA-directed DNA polymerase, pol protein, non-LTR retrotransposon, mobile element, retroelement, polyprotein, retrovirus and polymerase (access date Jan-2020).

To reduce ascertainment bias introduced by comparison to published elements, we complemented this final set with novel repeat sequences obtained from *LTR_Finder* 1.0.7 (Xu and Wang 2007), RepeatMasker 4.0.8 (Smit et al. 2013), and EDTA 1.9.9 (Ou et al. 2019). Identified sequences were pooled and used as reference in a second round of the whole pipeline, aiming to extend the finding of repeats that may differ from the already known reference sequences. Annotations of combined sequences retrieved from all packages were merged based on overlapping coordinates using BEDTools (Quinlan and Hall 2010). Additional to LTR elements, in the reference genome EDTA inferred 16 novel sequences belonging to TIR and non-TIR elements, not reported for *S. pombe* previously. 15 of these, however, overlapped with annotated functional genes suggesting false positive inference.

Different strains are known to have large structural variants, including inversions exceeding 1 Mbp in length and inter-chromosomal translocations (Brown et al. 2011; Teresa Avelar et al. 2013; Zanders et al. 2014; Tusso et al. 2019). In order to establish synteny between samples, all annotated coordinates were translated to the reference genome. For this, we produced a liftover of all genomic positions between each *de-novo* genome and the repeat-masked reference genome using flo (Pracana et al. 2017) and liftOver 2017-03-14 (Kent et al. 2002) requiring a minimum match of 0.7. Then we used customized Python and R scripts (R Core Team 2021) to identify translated coordinates of flanking sequences to breaking points for each TE element. Since TEs could occur in tandem several sequences will share the same adjacent, non-repetitive sequences within the sequence space of the masked reference genome and were grouped as clusters. In the case of several TE sequences per cluster, the position of the breaking points in the original de-novo genome were then shifted along the corresponding 3’ and/or 5’ axis until finding the first non-repetitive base in the lift over (**Supplementary Figure 15**). As a result, TE sequences within a cluster will all share the same flanking insertion breakpoint coordinates of the cluster. The final list of transposable sequences, the position of the cluster they belong to and their individual location and direction can be found in **Supplementary table 3**.

We compared the list of TE elements extracted in previous work to validate our pipeline in two ways. First, we compared LTRs detected in the reference genome by Bowen *et al*. (Bowen 2003) and our annotation of TE sequences in the reference genome; Second, we compared presence/absence scores from our data with scores based on paired-end short-read Illumina data from Jeffares *et al*. 2015 (**Supplementary table 4**). In the first case, we converted the cosmid-based coordinates of the LTRs annotated by Bowen et al. to coordinates in the current version of the reference genome by BLAST and manual adjustment (**Supplementary table 5**), and group sequences in clusters as we did for long read assemblies. We counted the number of sequences per cluster. For the second comparison in other samples and Jeffares *et al*. data set, we restricted this comparison to samples showing consistent strain ID between short- and long-read data. Differences observed between short- and long-read data (**Supplementary Figure 7**), were contrasted with differences observed between clonal strains using only *de-novo* genomes from long reads (**Supplementary Figure 6**).

### Phylogenetic analyses

We use a customized Python script to extract TE sequences of minimum length 100 bp from *de-novo* genomes. Consensus sequences from all samples produced were then used as reference for each query sequence. Query sequences were differentiated between solo-LTRs, fragmented TEs with one or no flanking LTR sequence, and full-length elements containing longer than 4.5 kb, the full polyprotein sequence and both flanking LTRs. Two alignments

were produced using MAFFT 7.407 (Katoh and Standley 2013): one for full-length TE; and another one for LTR sequences including solo-LTRs and all flanking LTRs associated to full-length TE elements. These alignments were used to produce a maximum-likelihood tree with IQ-TREE 1.6.10-omp-mpi (Nguyen et al. 2015) using the incorporated model prediction with ModelFinder (Kalyaanamoorthy et al. 2017) and 1000 ultrafast bootstrap (UFBoot) (Minh et al. 2013). Since different LTR families have been previously identified in the reference genome (Bowen 2003), we used genomic coordinates of known LTR sequences to place solo-LTR families for other strains in the phylogeny.

### Recombinant TE haplotypes

In order to identify potential recombinant haplotypes, the alignment of full-length elements was divided into windows of 30 bp. Other windows sizes like 20 and 10 bp were also tested with similar results. For each window, pairwise comparisons between sequences were performed. If sequences within windows differed by more than 2 bp, they were classified as different. When a sequence was equidistant to two already identified haplotypes, it was grouped to the first comparison. However, these cases were rather rare and do not have major impact on the general results. Then, to identify whole sequence haplotypes (all windows), pairwise comparisons between whole sequences was performed (**Supplementary Figure 16**). Haplotypes were scored as identical if they contained the same succession of identical 30 bp windows. We allowed one window to be different between sequences. Haplotypes were filtered, considering only haplotypes with at least 50% of the entire sequence and with at least 5 sequences (either paralogs or orthologs). This reduced the data set to 11 common haplotypes.

A similar analysis was performed for solo-LTRs and flanking LTRs. LTRs fragment were short (∼350 bp or ∼1060bp in the alignment including insertions and deletions) precluding the windows-based approach. Instead, we used the full alignments focusing on the most common LTR families (α and β). We identified potential recombinant haplotypes by looking first for diagnostic variants of each family. For this, diagnostic variants constituted those near-fixed between families (> 0.8 frequency in one family; <0.2 in the other) (**Supplementary Figure 9**). These diagnostic variants were contrasted with sequences in the Sub-α/ζ group and used to identify recombinant haplotypes. Sequences were grouped into haplotypes on the basis of pairwise comparisons of diagnostic variants. Two sequences were considered from a different haplotype if they differed in up to two diagnostic variants.

### Testing the ‘genomic shock hypothesis’

To test the hypothesis of TE reactivation in admixed genomes, we performed linear models to assess the relationship between admixture proportions (from pure *Sp* or *Sk* to 0.5 admixture) as explanatory variable and number of clusters containing at least one full-length TE as response. The normality assumption of the residuals held as assessed by the R package olsrr 0.5.3 setting a maximum p-value threshold of 0.05 (https://olsrr.rsquaredacademy.com/). Additionally, we restricted the analyses to the presumably more recent insertions including only those clusters consisting of a single full-length element in a single sample (**Supplementary Figure 13**). Analyses were performed including both TF1 and TF2 haplotypes, as well as by haplotype independently.

### Population genetic analyses

The frequency distribution of syntenic TE sequences among non-clonal strains was summarized in one- and two-dimensional site frequency spectra (SFS). Clusters sharing the same start and end coordinates, allowing for a 100 bp error margin on each side of the repetitive cluster, were defined as syntenic loci. Error margins of 150 and 200 bp yielded similar results. The error margin was necessary to account for variation introduced by the liftover. Spacing between clusters exceeded a minimum of 500 bp in all cases to guard against false positive inference of synteny of adjacent clusters (**Supplementary Figure 17**). Translating coordinates from de-novo genomes onto the reference genome could lead to a potential underestimation in the number of clusters. In cases where flanking sequences are absent in the reference genome or overlap with longer stretches of repetitive sequences, the range of a cluster will be expanded until the next non-repetitive anchor in the reference genome is found. As a consequence, non-orthologous, neighboring sequences of multiple query strains may be collapsed into a single genomic cluster spanning the ‘difficult-to-align region’ of the reference genome. However, even if this bias exists, it would be rare and will not affect the main conclusions because: i) TE insertions were distributed along the whole genome without gravitating towards repetitive regions (**Supplementary Figure 18**); ii) Tf1 insertions have been shown to be preferentially integrated into nucleosome-free promoters of genes (Bowen et al. 2003; Guo and Levin 2010); iii) the vast majority of clusters contained a single sequence per sample (**Supplementary Figure 3c,d**). Presence / absence of orthologues clusters was then scored as allelic state of the locus. Allele frequencies were summarized for all clusters by the derived state and summarized in an unfolded one-dimensional site frequency spectrum (**Figure 4a**). Only non-clonal strains were included to produce the SFS.

In addition, we obtained a two-dimensional site frequency spectrum (2dSFS) considering allele-frequency sharing between syntenic clusters surrounded by either *Sp* and *Sk* ancestry. Ancestry was inferred by classifying SNP information in the flanking sequences of insertion breakpoints for each cluster and sample (Tusso et al. 2019). For each locus, allele frequencies were separately estimated for each ancestral background resulting in an unfolded 2dSFS. In addition to simple presence/absence scoring, alleles were scored according to the number of sequences within cluster, their family and direction of insertion – yielding comparable results (**Figure 4b**, **Supplementary Figure 19, and Supplementary table 6**). In both cases, only variants with at least 4 samples per genetic background were considered.

One- and two-dimensional site frequency spectra were calculated for all TE sequences (**Figure 4B**) and separately for full-length elements (**Figure 4C**). A two-dimensional site frequency spectrum was likewise produced from non-coding genome-wide SNPs using the same set of non-clonal strains, and removing repetitive regions. This resulted in a final data set of 209,690 variant sites. Allele frequencies to produce a folded two-dimensional site frequency spectrum were calculated using the R package SNPRelate 1.24.0 (Zheng et al. 2012). In the absence of an appropriate outgroup, SNP variants cannot be polarized into an ancestral and derive state. To allow direct comparisons between SFS between TEs and SNPs, the two-dimensional site frequency spectrum of TEs was also folded (**Supplementary Figure 14**).

### Sequence divergence of LTRs

To assess the levels of sequence divergence of TEs, we calculated divergence between all solo-LTR and flanking LTR sequences as a function of the genomic background (*Sp* or *Sk*) they are embedded in. We divided sequences by cluster and family according to phylogenetic reconstruction (**Figure 2A**), and used the R package ape 5.4-1 (Paradis and Schliep 2019) to calculate pair-wise Kimura’s 2-parameters distance (Kimura 1980) within and between ancestral backgrounds. Additionally, we measured divergence between the 5’ and 3’ flanking LTR sequences of each full-length TE element as a proxy of the age of its individual insertion (Supplementary Figure 1**2**).

## Data Access

The PacBio sequencing data and genome assemblies generated in this study have been submitted to the CNGB Sequence Archive (CNSA) (https://db.cngb.org/cnsa) under Project ID CNP0001878. All code used for the analyses are available in Supplemental Code and at zenodo DOI: 10.5281/zenodo.5747176.

## Acknowledgments

We thank S. Lorena Ament-Velásquez, Fidel Botero-Castro, Bart P.S. Nieuwenhuis, Claire Peart, Ricardo Pereira, Alexander Suh, Vera Warmuth, Matthias Weissensteiner and members of the Wolf and Du labs for providing intellectual input on the various analyses, and comments on the manuscript. Funding was provided by LMU Munich (JW). The computational infrastructure was provided by the UPPMAX Sequencing Cluster and Storage (UPPNEX) project funded by the Knut and Alice Wallenberg Foundation and the Swedish National Infrastructure for Computing.

## Contributions

ST, LLD and JW conceived the study; All analyses were performed by ST and FS with contributions from YL in genome annotation of transposable elements analyses. ST and JW wrote the manuscript with input from FS, YL and LLD.

