## Supplementary Figures for "Reactivation of transposable elements following hybridization in fission yeast"

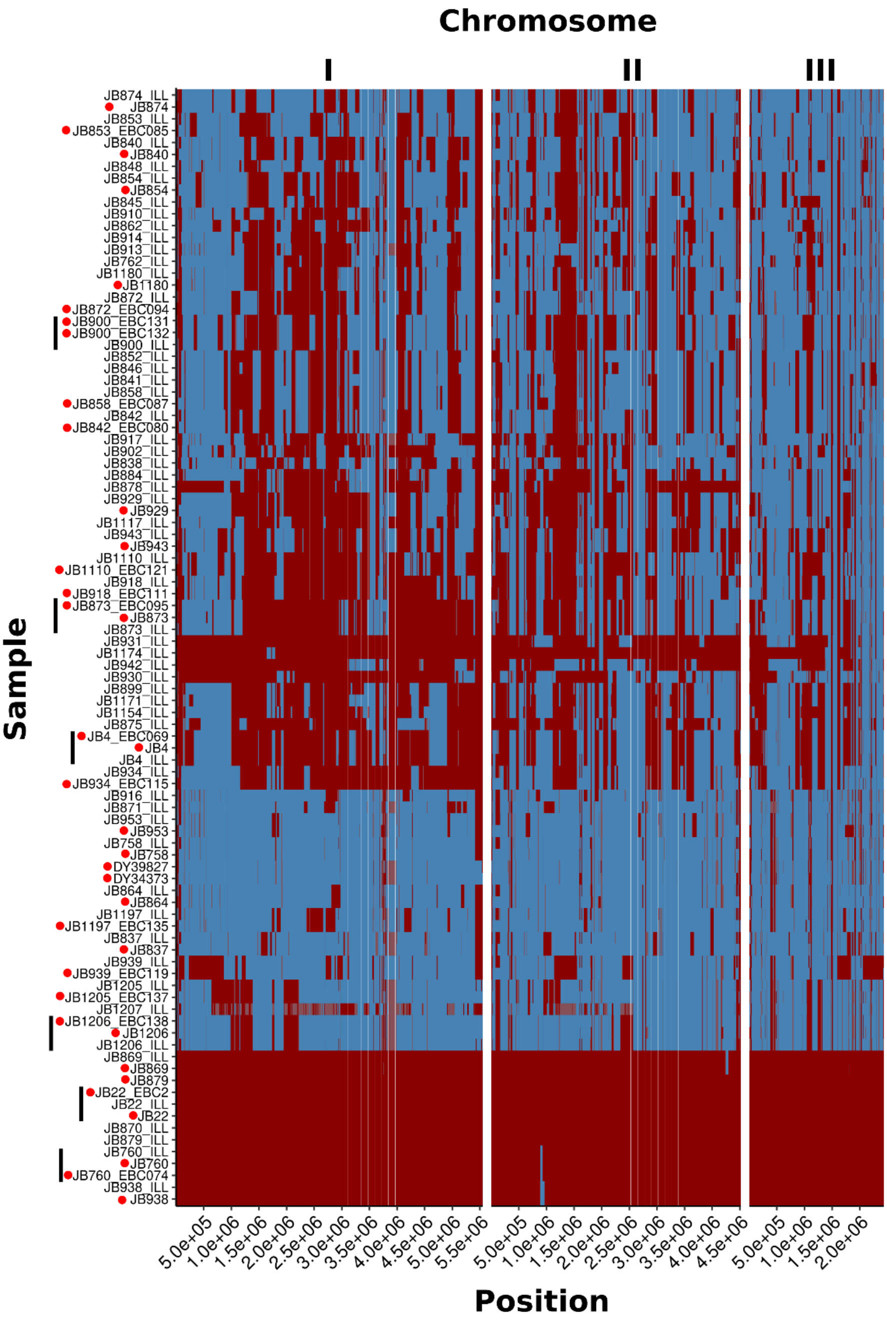


**Supplementary Figure 1**: Heatmap representing SNP-based haplotypes across all three chromosomes for the 57 strains representing global diversity of *S. pombe* (Jeffares et al., 2015). The 37 strains for which long-read based de novo assemblies have been generated for this study are indicated by red dots. Six instances of clonal samples derived from the same strains are indicated by an additional black bar. Haplotypes are painted by *Sp* and *Sk* ancestry indicated by red and blue colour, respectively (Tusso et al., 2019). The suffix ILL of strains ID reflects that ancestry inference was based on Illumina short reads. For all other strains, ancestry was inferred on the basis of long reads from single-molecule real-time sequencing. Note that ancestry inference is independent of the sequencing method.

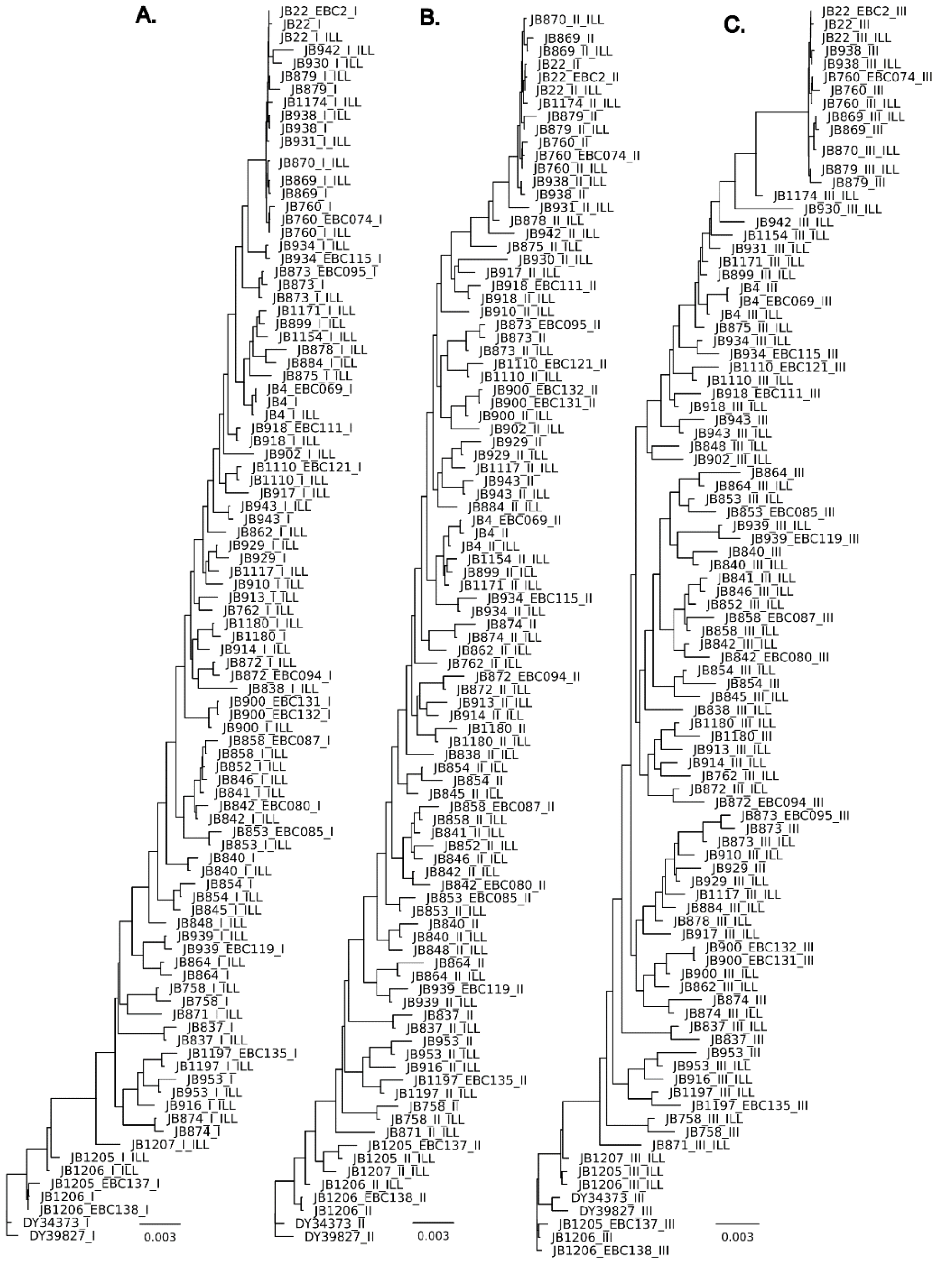


**Supplementary Figure 2:** Un-rooted maximum likelihood trees for both short- and long-read data. IDs of strains sequenced only with short-read data are labeled with *ILL*. Panels **A**, **B** and **C** represent phylogenies for chromosome I, II, and III respectively.

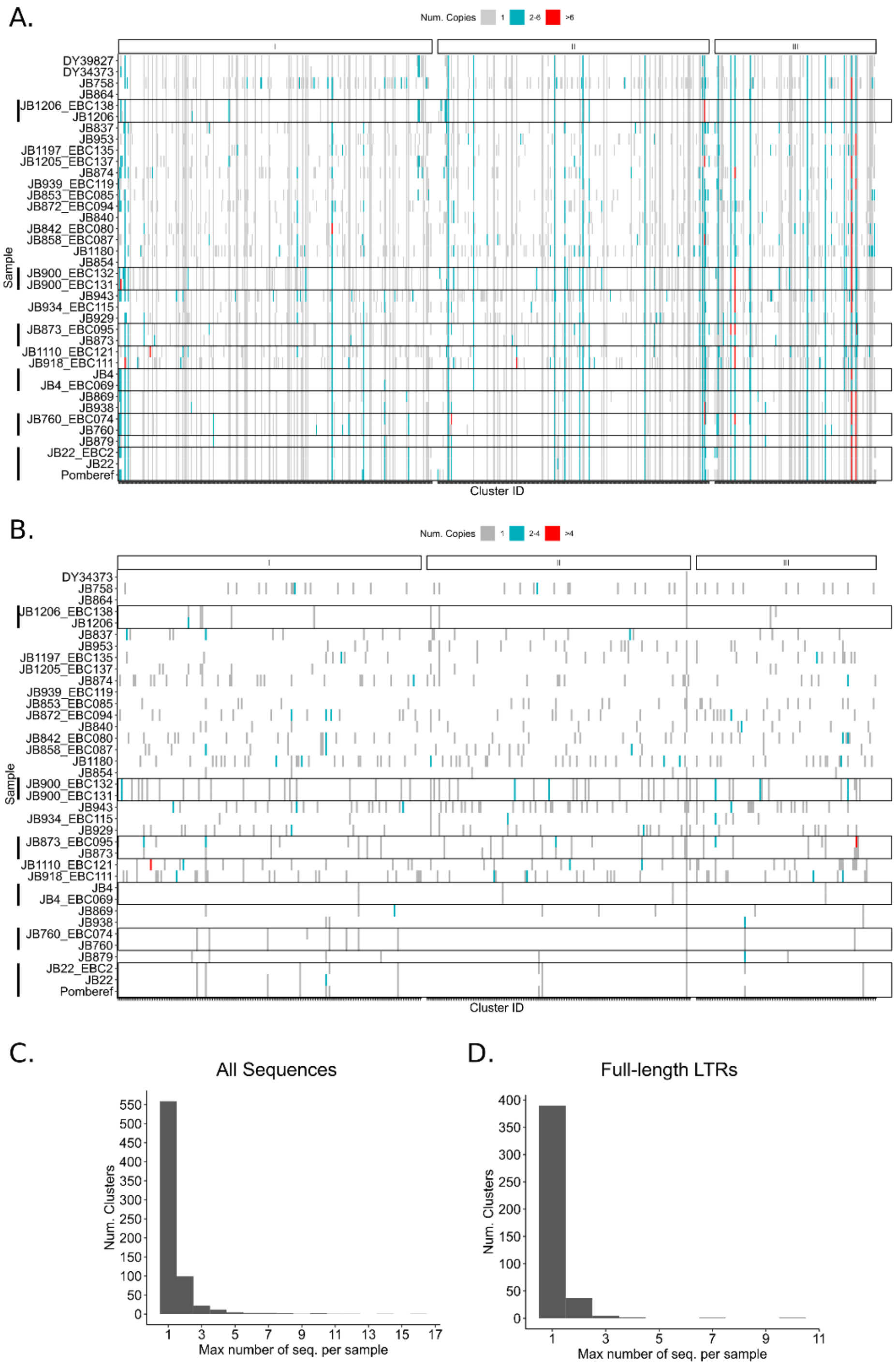


**Supplementary Figure 3:** Heatmap displaying syntenic TE clusters along the genome for all 38 samples used in this study. Colours indicate the number of TE sequences inserted per cluster in three categories range. **A.** Plot with all sequences, **B.** plot restricted to full-length sequences. Clonal samples from the same strain are grouped with a black box and indicated with black bars. Strains are ordered according to ancestral proportion, from pure *Sk* on top to pure *Sp* strains at the bottom. **C.** histogram of the number of clusters with a given maximum number of sequences per sample. **D.** As in c, but only including full-length LTR elements.

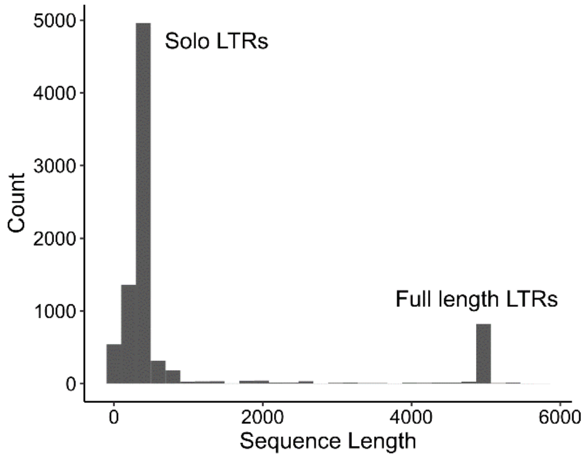


**Supplementary Figure 4:** Histogram of the length distribution of all TE elements found in the global collection of 38 samples derived from 31 *S. pombe* strains.

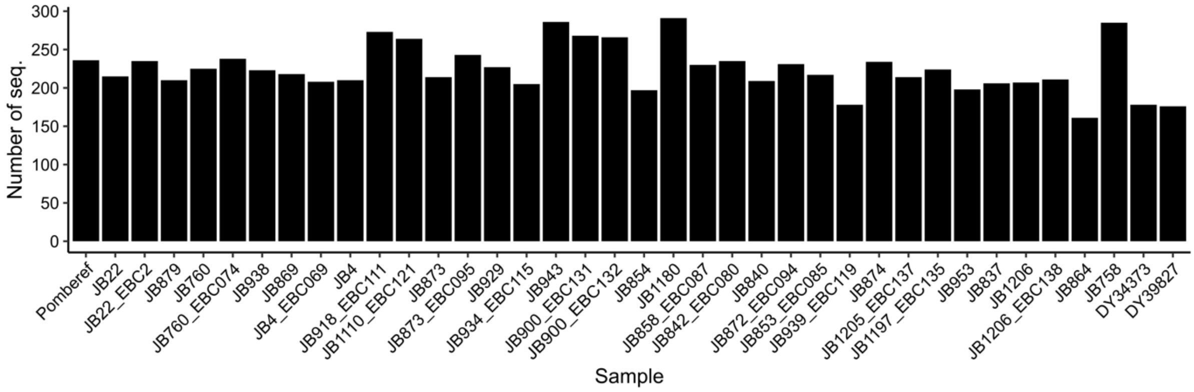


**Supplementary Figure 5:** Total number of TE sequences per sample including solo-LTRs as well as full-length elements.

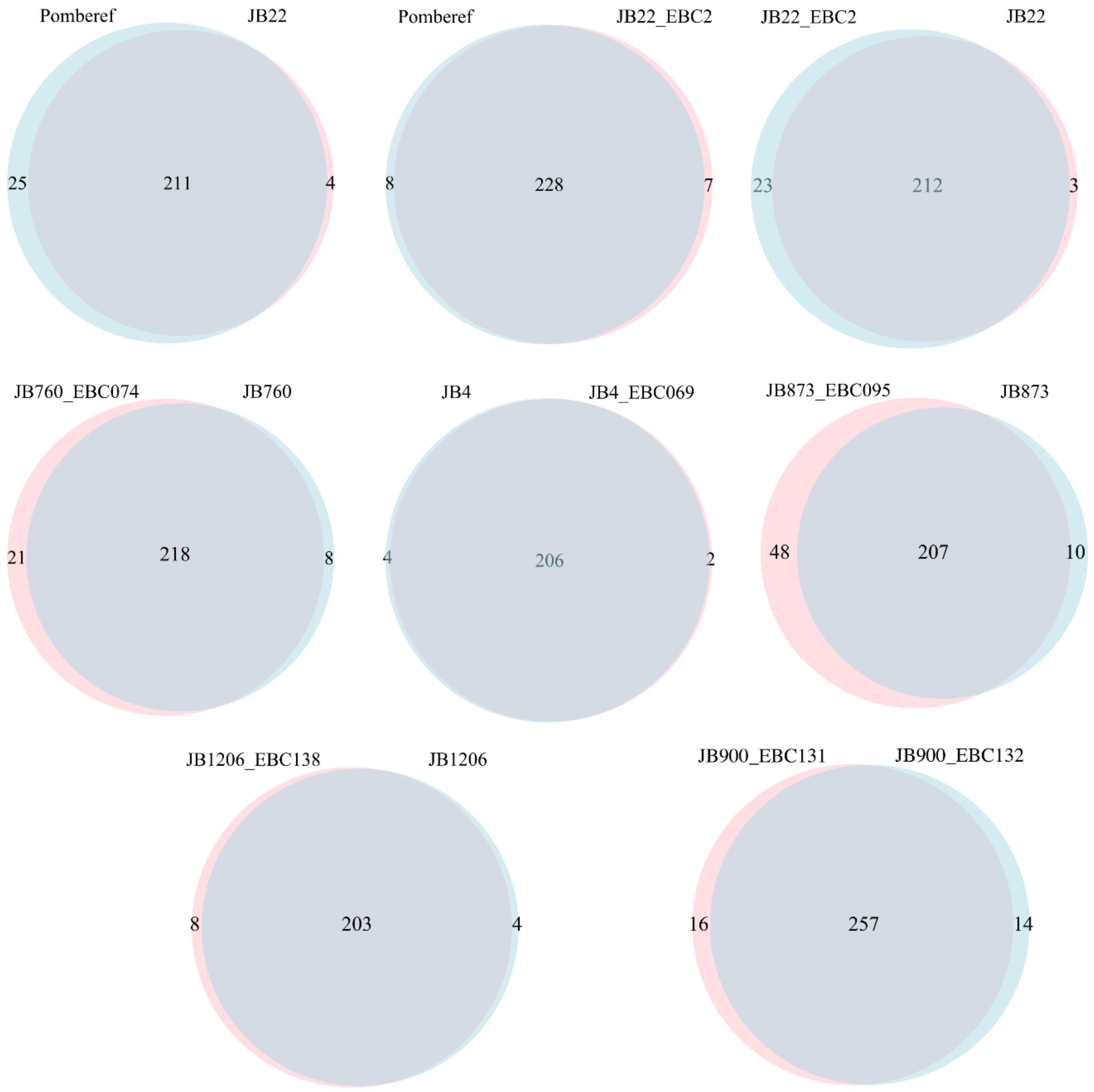


**Supplementary Figure 6:** Venn-diagrams displaying pair-wise comparisons between clonal samples derived from the same 6 strains (JB22, JB760, JB4, JB873, JB1206, JB900). For each sample comparison, the diagram quantifies the shared and unique number of sequences.

For example, consider comparisons among the *S. pombe* reference genome (ID: Pomberef) and *de novo* genomes derived from the same strain (IDs: JB22_EBC2 and JB22) revealing a number of differences. In Pomberef, we identified a total of 189 clusters, containing 236 TE sequences. In contrast, in JB22_EBC2 and JB22_DY38751, we scored 185 and 175 clusters with 236 and 215 TE sequences, respectively. While all 185 clusters from JB22_EBC2 were shared with Pomberef, two clusters, each containing a single TE sequence, were specific to JB22. Some differences corresponded to new sequence insertions present only in either JB22_EBC2 or JB22, or conversion from full-length elements to solo-LTRs.

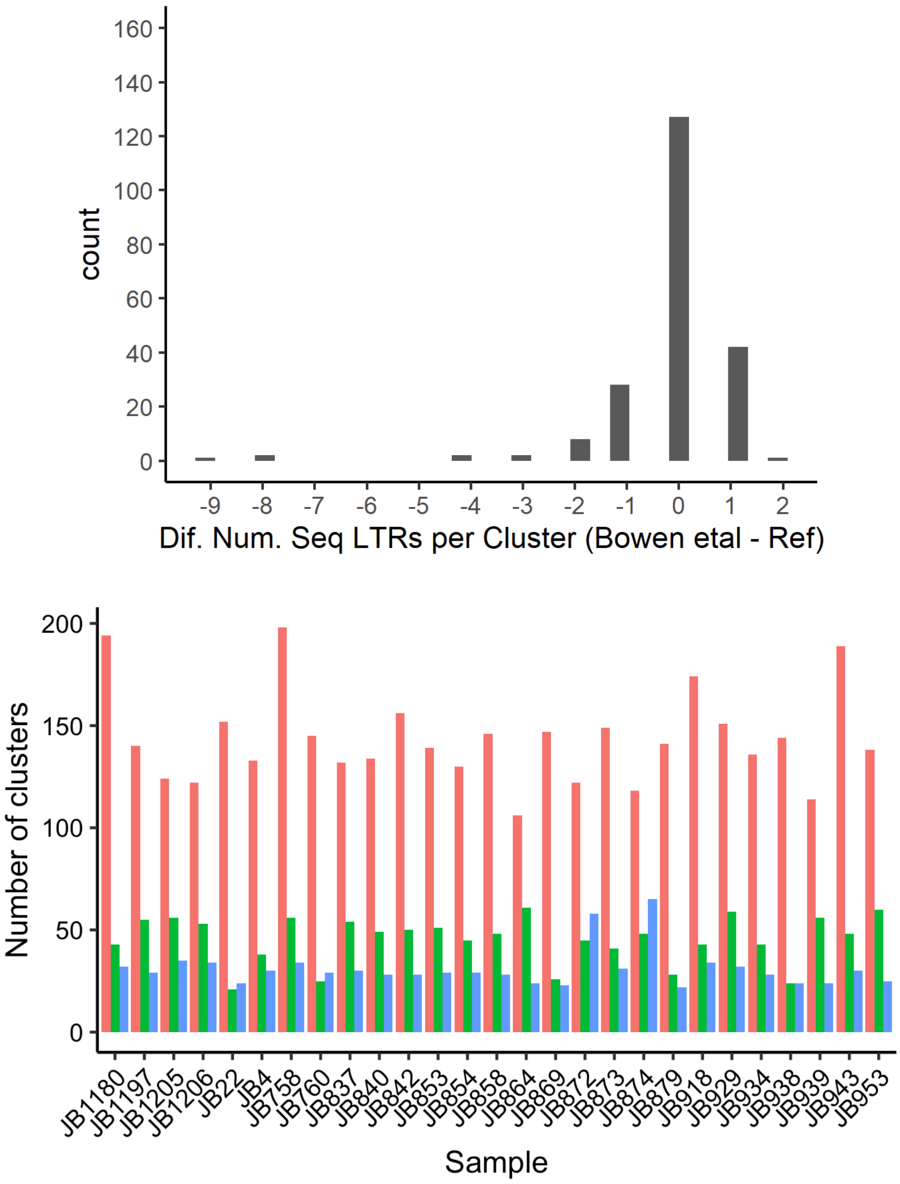


**Supplementary Figure 7:** Comparison of TE sequences found in this study and previous studies based on short-read data. **top panel.** Comparison of the reference genome and sequences reported in Bowen *et al.* 2013. The histogram shows the difference in the number of TE elements (both solo-LTR and full-length) within each syntenic cluster. **bottom panel.** Comparison with TE calls from Jeffares *et al.* (2015) for 27 of the 31 strains used in this study. In this data set, sequences within syntenic clusters are not reported, thus for each strain, the number of clusters is reported. The three bars show orthologous clusters found in both data sets (red), clusters found exclusively by Jeffares et al. (2015) (green) and clusters only identified in this study (blue).

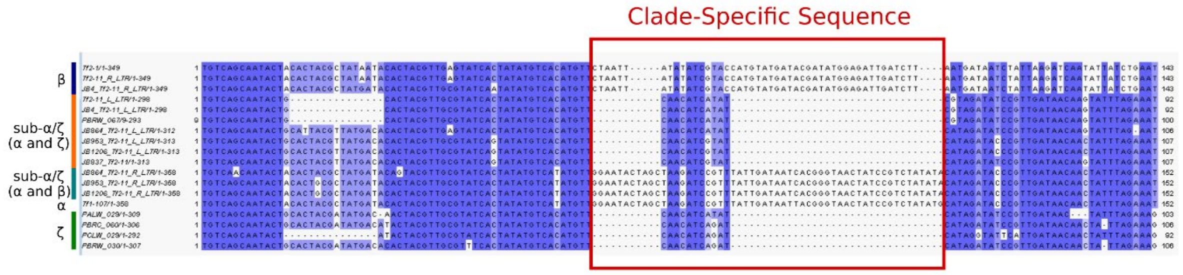


**Supplementary Figure 8:** Alignment illustrating the similarity between ζ LTR and first half of the sub-α/ζ LTR. The alignment contains flanking LTR from a Tf2 (β) and Tf1(α) haplotype as comparison. The red box highlights a clade-specific sequence which is deleted in the ζ and sub-α/ζ families.

A.

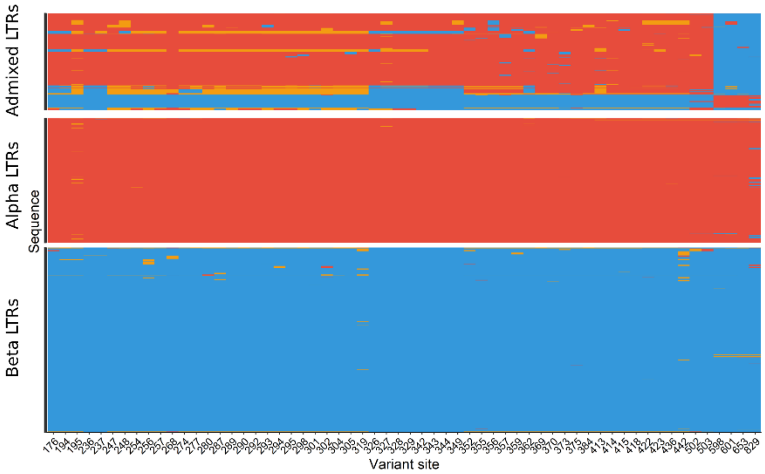


B.

| ID | All Seq. | | | Full-LTRs | | |
|---|---|---|---|---|---|---|
| | Seq. | Sam. | Clus. | Seq. | Sam. | Clus. |
| 2 | 523 | 38 | 64 | 72 | 19 | 31 |
| 659 | 110 | 20 | 40 | 96 | 18 | 29 |
| 573 | 12 | 12 | 3 | | | |
| 369 | 34 | 27 | 2 | 19 | 16 | 2 |
| 514 | 2 | 2 | 2 | 1 | 1 | 1 |
| 574 | 18 | 18 | 2 | 18 | 18 | 2 |
| 414 | 5 | 5 | 1 | 5 | 5 | 1 |
| 416 | 4 | 4 | 1 | 3 | 3 | 1 |
| 552 | 21 | 21 | 1 | | | |
| 575 | 29 | 29 | 1 | 2 | 2 | 1 |
| 593 | 1 | 1 | 1 | 1 | 1 | 1 |
| 594 | 26 | 25 | 1 | | | |
| 621 | 15 | 14 | 1 | | | |
| 653 | 1 | 1 | 1 | | | |
| 770 | 1 | 1 | 1 | | | |
| 771 | 4 | 4 | 1 | | | |
| 775 | 27 | 27 | 1 | | | |
| 802 | 1 | 1 | 1 | | | |
| 803 | 37 | 37 | 1 | | | |
| 870 | 8 | 8 | 1 | | | |
| 878 | 1 | 1 | 1 | | | |
| 879 | 1 | 1 | 1 | 1 | 1 | 1 |
| 880 | 1 | 1 | 1 | 1 | 1 | 1 |
| 881 | 1 | 1 | 1 | | | |

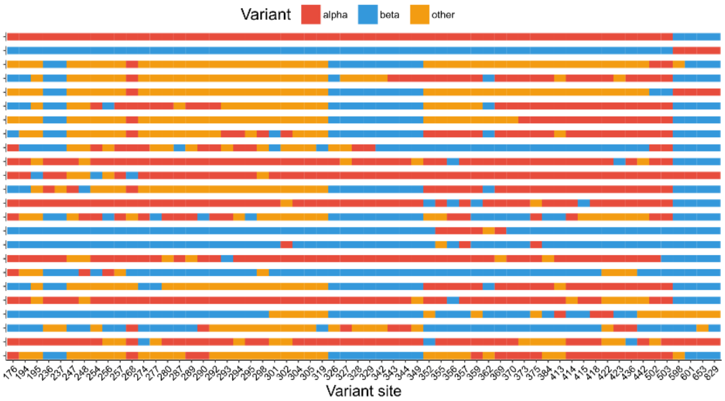


C.

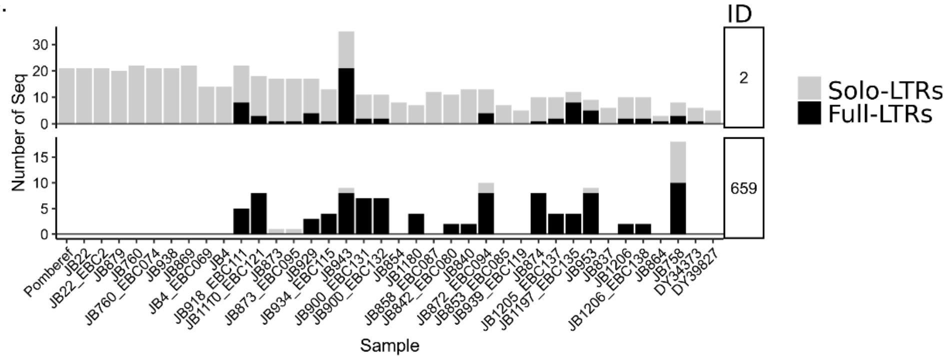


**Supplementary Figure 9**: Insight into recombinant TE haplotypes. **A.** Comparison between alpha, beta and recombinant LTR haplotypes (both solo-LTR and flanking LTRs for full length elements). The figure shows diagnostic variant sites for the alpha and beta haplotypes. Positions correspond to a location of the total alignment spanning 1,063 bp. Diagnostic variants were identified as the major allele frequency within group, considering only variants with major allele frequency higher than 0.8, but lower than 0.2 in the other group. **B.** Summary of admixed LTR haplotypes. For each haplotype we tabulated the total number of instances the haplotype occurred across all samples (Seq.), the number of samples containing the haplotype (Sam.) and the number of clusters within which it occurred (Clus.). Values differentiate between all LTR sequences and restricted to flanking LTRs from full-length elements. Haplotypes found in locus Tf2-11 are highlighted in grey. **C.** Bar plot showing the number of sequences per sample for the two most common recombinant haplotypes (IDs: 2, 659). For each sample, the number of solo-LTRs and flanking LTRs of full-length elements are shown. Samples are organized from ‘pure’ *Sp* ancestry to ‘pure’ *Sk* ancestry.

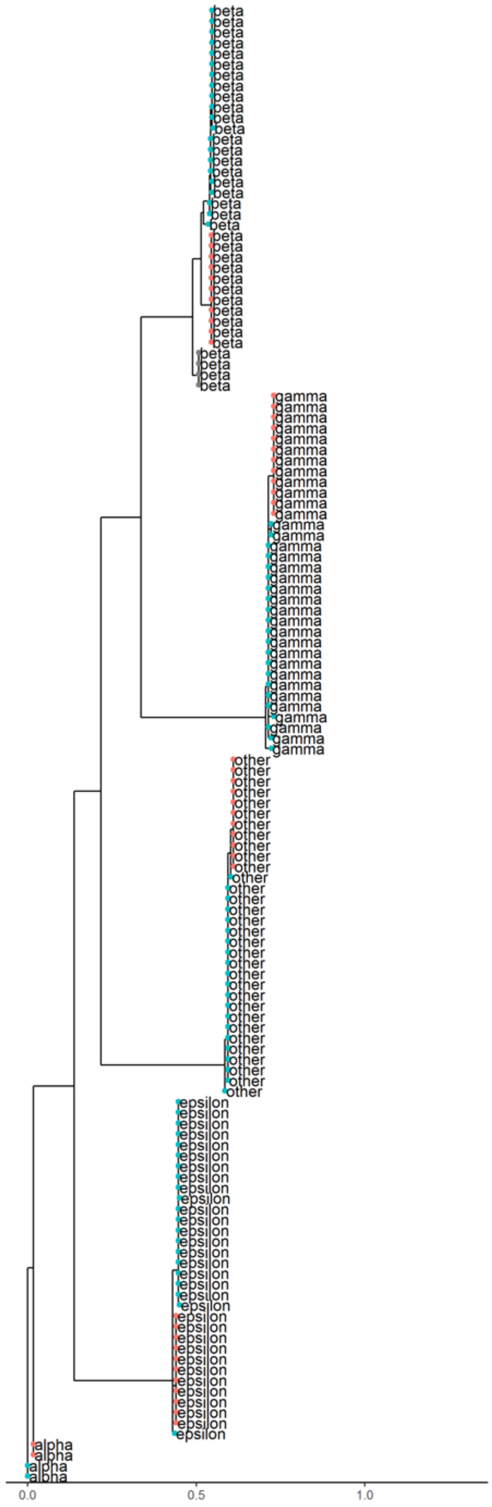


**Supplementary Figure 10:** Example of a maximum likelihood tree of LTR sequences from within a single TE cluster, illustrating large-scale divergence between LTR families, and shallow differentiation of LTRs from the same family between the *Sp* (red) and *Sk* (blue) ancestral background. This example corresponds to cluster 582 in chromosome II, start position 4436600. Points show ancestral group in flanking sequences (red:*Sp*, Blue:*Sk*). Labels indicate LTR family in **Figure 2.**

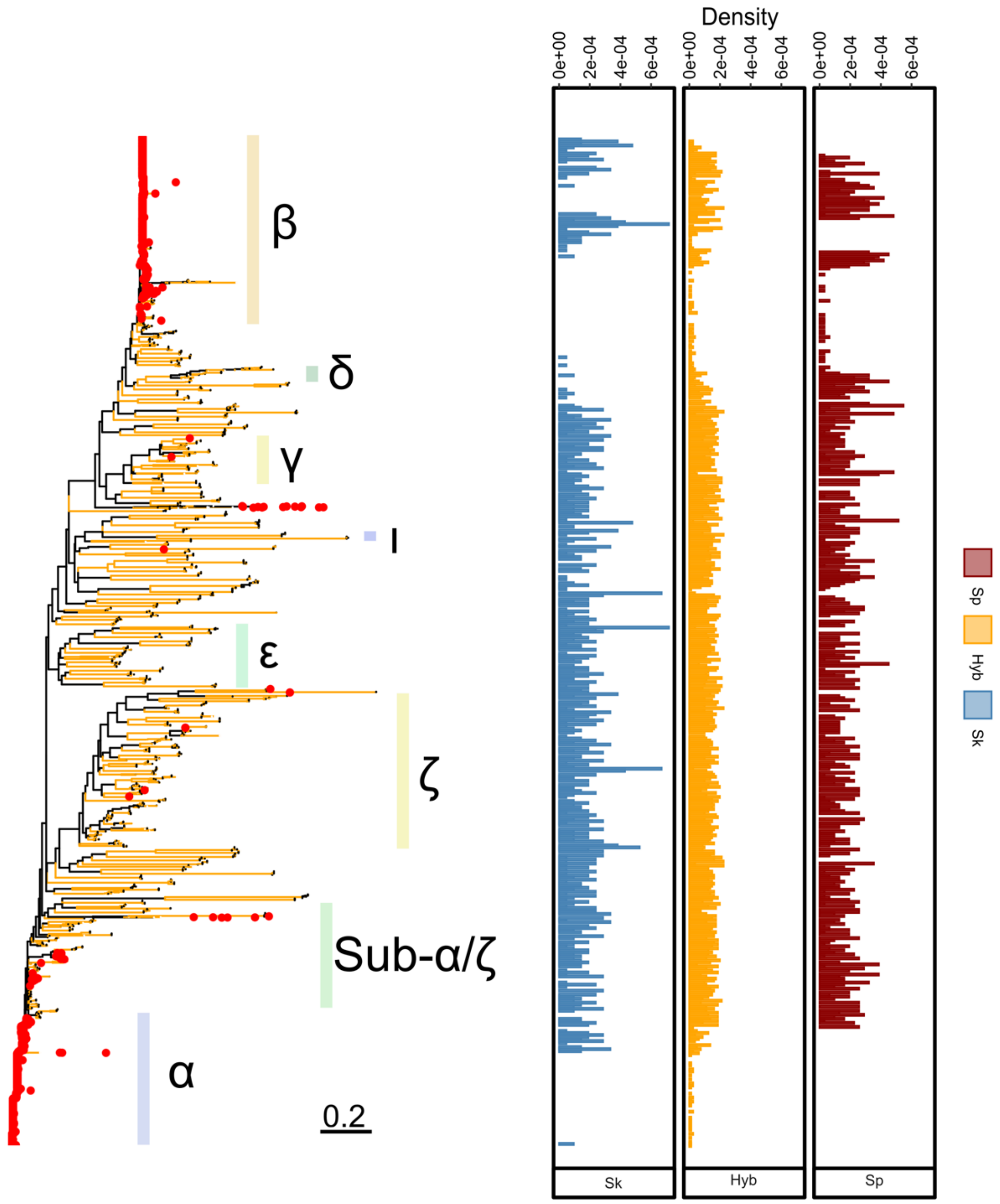


**Supplementary Figure 11:** Maximum likelihood tree of LTR sequences as in **Figure 2a**. Right panels show density histograms of abundance of solo-LTRs with rows corresponding to the respective branch on the phylogeny for non-admixed samples (Sk: blue or Sp: red) and hybrid samples (Hyb: yellow).

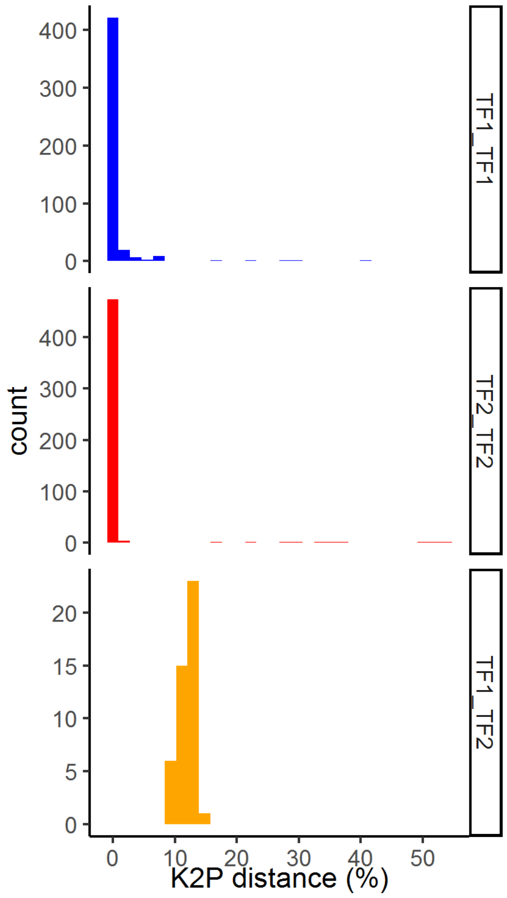


**Supplementary Figure 12:** Pair-wise divergence between flanking LTRs contained in full-length elements. Sequences were divided between those without signatures of recombination in **Figure 3** (here shown as either Tf1_Tf1 or Tf2_Tf2), and those with recombinant haplotypes (Tf1_Tf2). Divergence was measured as Kimura's 2-parameters distance (Kimura, 1980).

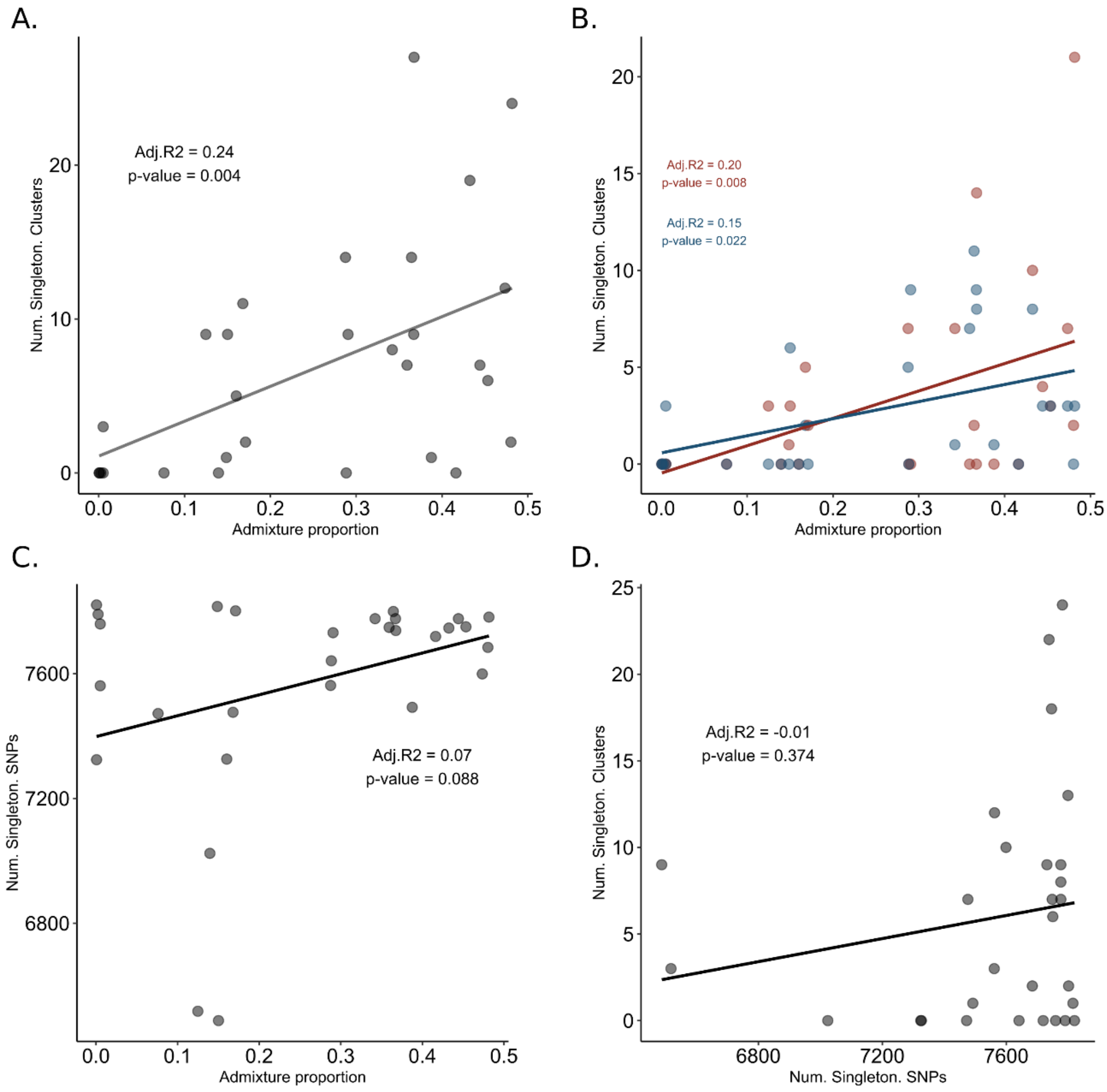


**Supplementary Figure 13:** Relationship between genomic ancestry and TE abundance. Correlation between ancestral admixture proportion and the number of singleton clusters containing a full-length element. The analysis was done including both Tf1 and Tf2 haplotypes (**A**) or independently (**B**). **C.** Correlation between ancestral admixture proportion and the number of genome-wide singleton SNPs. Only SNPs in non-coding regions were included. **D.** Correlation between number of genome-wide singleton SNPs and number of singleton clusters containing a full-length element. Each point represents one non-clonal strain. Adjusted $R^2$ and p-value of the linear model are shown as inset.

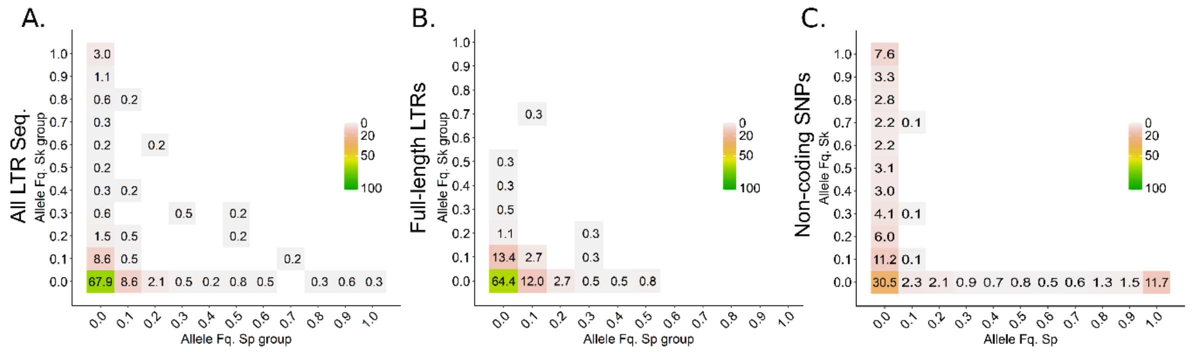


**Supplementary Figure 14:** Folded two-dimensional site frequency spectrum illustrating shared allele frequencies between ancestral *Sp* and *Sk* backgrounds for (**A**) all LTR sequences, (**B**) full-length elements and (**C**) non-coding genome-wide SNPs. Numbers and colour scale show the proportion of variants falling into each allele frequency bin.

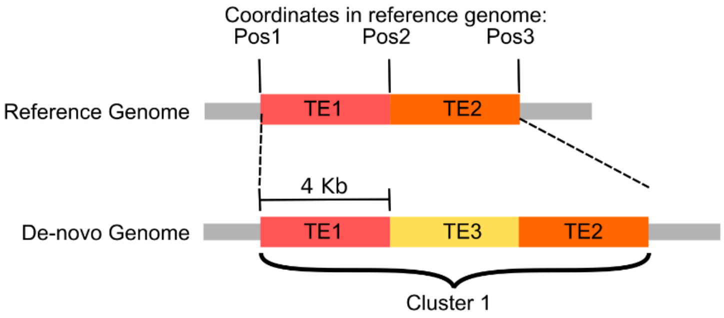


Output:

| | Start pos in reference | Shift start pos | End pos in reference | Shift end pos | Cluster |
|---|---|---|---|---|---|
| TE1 | Pos1 | 0 | Pos3 | 8kb | 1 |
| TE2 | Pos1 | 8kb | Pos3 | 0 | 1 |
| TE3 | Pos1 | 4kb | Pos3 | 4kb | 1 |

**Supplementary Figure 15:** Example of liftover output translating coordinates of TE elements (solo-LTRs and full-length elements) within clusters between the *S. pombe* reference genome and the *de-novo* assemblies from this study. Overlapping TE sequences, or sequences arranged in tandem are considered to belong to the same syntenic genomic cluster. Inferred start and end point for sequences within the cluster are the same, but the distance to the breaking points is recorded.

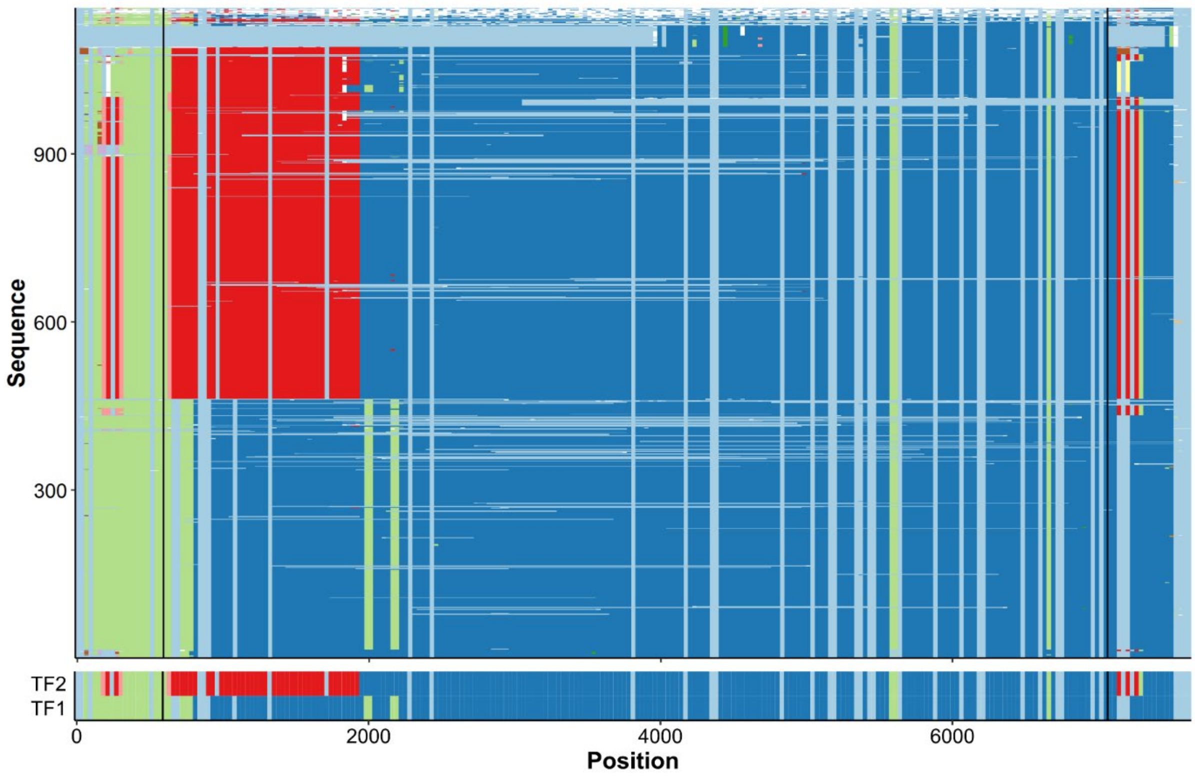


**Supplementary Figure 16:** Alignment of the all, unfiltered haplotypes of full-length LTR elements identified by window-based haplotype painting in a global sample of *S. pombe*. Window colours represent difference haplotypes within windows (by columns). Gaps in the alignment or in incomplete sequences are shown in light blue. As reference haplotype Tf1 and Tf2 are shown in the bottom.

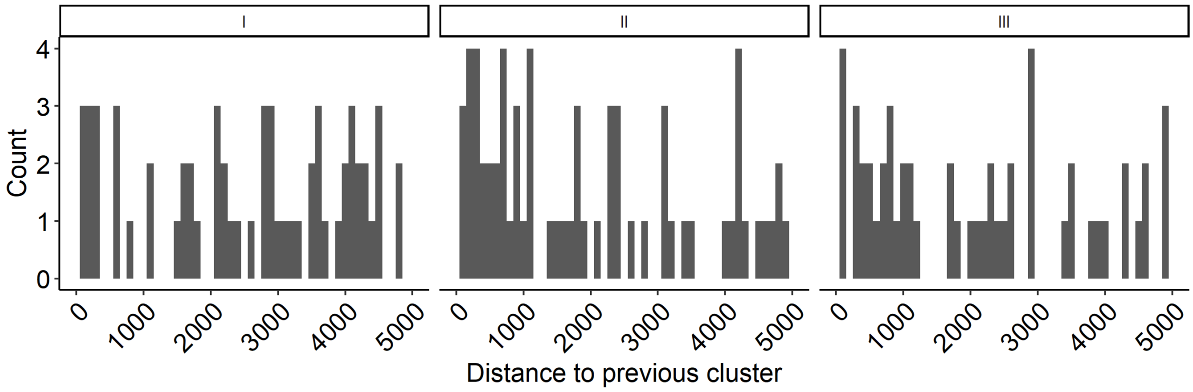


**Supplementary Figure 17:** Histogram tabulating the number of clusters falling within a given distance [bp] to the previous cluster. Panels are divided by chromosome.

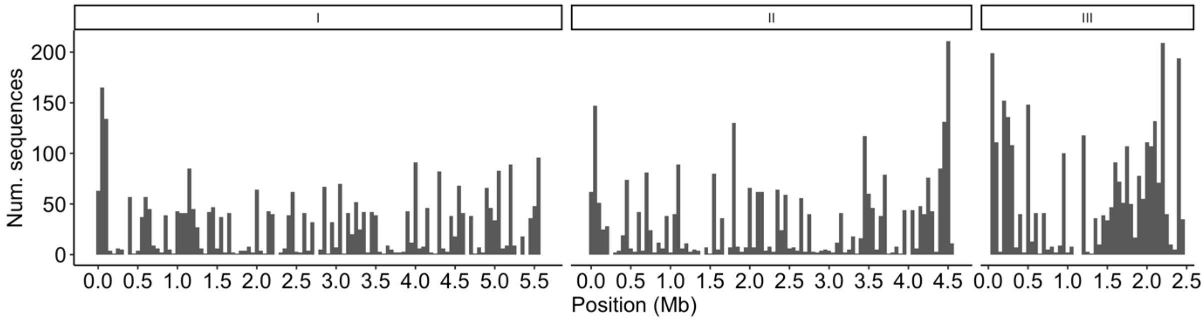


**Supplementary Figure 18:** Histogram tabulating the number of TE sequences along the genome combining all samples. Panels are divided by chromosome.

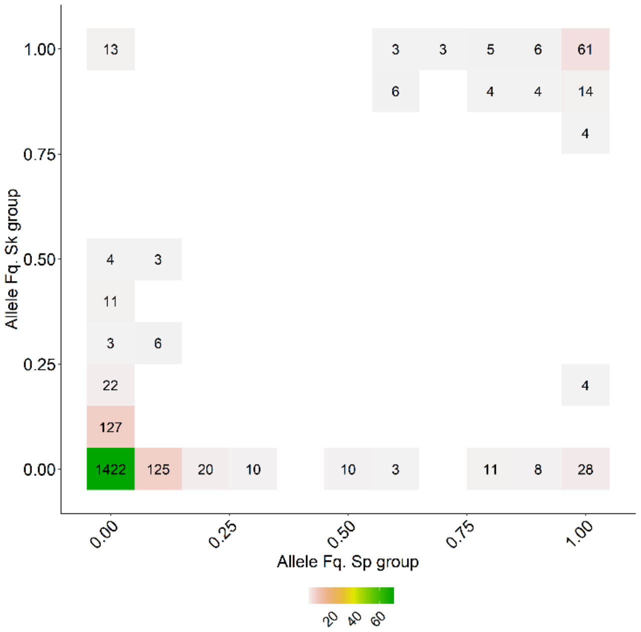


**Supplementary Figure 19:** Two-dimensional site frequency spectrum using solo-LTRs and flanking LTRs of full-length elements. Alleles are scored taken family identity and insertion direction into account. Frequencies thus represent the number of closely related LTRs per cluster. The colour scale indicates the proportion of sequences for each frequency bin.
